# Supervised spatial inference of dissociated single-cell data with SageNet

**DOI:** 10.1101/2022.04.14.488419

**Authors:** Elyas Heidari, Tim Lohoff, Richard C. V. Tyser, John C. Marioni, Mark D. Robinson, Shila Ghazanfar

## Abstract

Spatially-resolved transcriptomics uncovers patterns of gene expression at supercellular, cellular, or subcellular resolution, providing insights into spatially variable cellular functions, diffusible morphogens, and cell-cell interactions. However, for practical reasons, multiplexed single cell RNA-sequencing remains the most widely used technology for profiling transcriptomes of single cells, especially in the context of large-scale anatomical atlassing. Devising techniques to accurately predict the latent physical positions as well as the latent cell-cell proximities of such dissociated cells, represents an exciting and new challenge. Most of the current approaches rely on an ‘autocorrelation’ assumption, i.e., cells with similar transcriptomic profiles are located close to each other in physical space and vice versa. However, this is not always the case in native biological contexts due to complex morphological and functional patterning. To address this challenge, we developed SageNet, a graph neural network approach that spatially reconstructs dissociated single cell data using one or more spatial references. SageNet first estimates a gene-gene interaction network from a reference spatial dataset. This informs the structure of the graph on which the graph neural network is trained to predict the region of dissociated cells. Finally, SageNet produces a low-dimensional embedding of the query dataset, corresponding to the reconstructed spatial coordinates of the dissociated tissue. Furthermore, SageNet reveals spatially informative genes by extracting the most important features from the neural network model. We demonstrate the utility and robust performance of SageNet using molecule-resolved seqFISH and spot-based Spatial Transcriptomics reference datasets as well as dissociated single-cell data, across multiple biological contexts. SageNet is provided as an open-source python software package at https://github.com/MarioniLab/SageNet.

## INTRODUCTION

Spatial transcriptomics is an emerging technology (Marx 2021; Moses and Pachter 2021) that enables the study of intercellular networks, spatially associated subpopulations and transcriptional patterns, yielding a better understanding of intercellular communication and tissue patterning (Marx 2021), which plays a pivotal role in development (Barnes 2020; Han *et al.* 2020), cancer and disease progression (Saviano, Henderson, and Baumert 2020; Winkler *et al.* 2020), and homeostasis (Yang *et al.* 2019). Existing atlas-scale single-cell RNA-sequencing (scRNA-seq) compendia (Regev *et al.* 2017) have profiled the expression of millions of cells; however, their spatial context is lost. Thus, there is an exciting opportunity to recover the latent spatial dimension of such datasets.

Two key conditions need to be met in order to reconstruct the latent spatial dimensions of tissues captured from scRNA-seq data, independent of the presence of a spatial transcriptomic reference dataset. First, tissue niches should be characterizable by specific gene expression signatures. Second, these gene expression signatures should be reliably captured by the experimental platform. Thus, the ability to perform spatial reconstruction depends on the experimental data as well as the biological tissue of interest. Examples where such principles have been used include reconstruction of expression patterns along the major anatomical axes of the mammalian Organ of Corti from single cell qPCR (Waldhaus, Durruthy-Durruthy, and Heller 2015), and ordering of hepatocytes along a porto-central axis derived from scRNA-seq (Payen *et al.* 2021). However, for less well-characterised or complex tissue contexts, or at a whole-organism scale, the task of spatial reconstruction becomes more challenging, with a need for better methodological approaches that can complement spatial transcriptomics reference data.

Existing approaches to map dissociated single cells to a spatial coordinate system have ranged in input data requirements, underlying assumptions, as well as methodological focus. One general approach is to assume that gene expression profiles correspond to spatial coordinates, i.e., cells that are physically proximal are likely to exhibit a similar gene expression profile. Consequently, mapping single cells onto a spatial coordinate system is equivalent to performing appropriate dimensional reduction on the entire gene expression matrix or some meaningful subset of genes (Karaiskos *et al.* 2017, Nowotschin *et al.* 2019). Specific approaches for inferring the physical space have varied in complexity, from applications of classical dimensionality reduction techniques such as Principal Component Analysis (Durruthy-Durruthy *et al.* 2014, Mori *et al.* 2019, Ren *et al.* 2020) or latent variable models (Verma and Engelhardt 2020) to approaches using concepts from optimal transport, such as novoSpaRc (Moriel *et al.* 2021) and SpaOTsc (Cang and Nie 2020). In all cases, a key limitation of these approaches is the underlying assumption: cells with similar gene expression profiles are proximal in space. This assumption is not generally true, for example, endothelial cells with similar transcriptional profiles are present in multiple distinct locations across mouse embryos (Lohoff *et al.* 2021), or tumour tissues (Ali *et al.* 2020). Additionally, such approaches are unable to account for situations where cells with distinct gene expression profiles are proximal to each other in space, as happens in the layered structure of the mouse cortex (Maynard *et al.* 2021) and hippocampus (Eng *et al.* 2019), thereby precluding the ability to fully capture the biology of cell communication between distinct cell types (Han *et al.* 2020; Almet *et al.* 2021).

Given a spatially-resolved reference dataset, an additional class of approaches aim to directly predict the latent coordinates of tissue captured with scRNA-seq. For example, Tangram (Biancalani *et al.* 2020) uses deep learning to map dissociated cells directly to physical locations given a spatial reference, while other approaches (e.g., a convolutional neural network) have been tailored to predict the spatial origin of single neurons (Ortiz *et al.* 2020). These approaches may suffer from lack of robustness in generalizing to new unseen contexts, biological interpretability, and inability to provide measures of confidence, due to the nature of ‘black-box’ deep learning approaches (Hendrycks and Dietterich 2019; Abdar *et al.* 2021; Ching *et al.* 2018).

Here, we introduce SageNet, a method that reconstructs latent cell positions by probabilistically mapping cells from a dissociated scRNA-seq query dataset to non-overlapping partitions of a spatial molecular reference. SageNet uses the spatially-resolved transcriptomics reference to estimate a gene interaction network (GIN), which then forms the scaffold for training a graph neural network (GNN). The GIN provides the GNN an inductive bias to make the output more robust (Battaglia *et al.* 2018). SageNet outputs a probabilistic mapping of dissociated cells to spatial partitions, an estimated cell-cell spatial distance matrix, as well as a set of spatially informative genes (SIGs). From the reconstructed cell-cell distances, SageNet additionally outputs a low-dimensional embedding of cells for visualisation, and a cell-wise score representing the confidence of mapping. We demonstrate that SageNet is accurate, efficient, and robust using various datasets across technologies and biological contexts.

## RESULTS

### Spatial reconstruction by probabilistically assigning query cells to spatial partitions

SageNet probabilistically maps scRNA-seq query cells to non-overlapping partitions of a spatial reference using graph neural networks (GNNs). The input for training the model is one or multiple spatial reference dataset(s), where each is decomposed into non-overlapping partitions, either specified *a priori* or estimated by clustering based on the spatial coordinates (Figure 1, Supplementary Figure 1A). SageNet facilitates the use of multiple partitionings of the same spatial reference (e.g., at different resolutions) in order to capture spatially-associated gene expression patterns at multiple tissue scales. To train each GNN, a gene interaction network (GIN) is required, either input from the user or estimated from the reference data using the graphical LASSO (Figure 1, Supplementary Figure 1A, Methods, (Friedman, Hastie, and Tibshirani 2008)). The GIN is employed by the GNN as an inductive bias, representing the notion that genes interact with each other to form functional modules and that the spatial information can be encoded in this gene network. An ensemble SageNet model is built by combining the GNN models for each spatial reference dataset and partition (Methods). The most important features of a trained SageNet model can be extracted using the integrated gradients technique ((Sundararajan, Taly, and Yan 2017), Methods), allowing biological interpretation of such SIGs.

**Figure 1.**
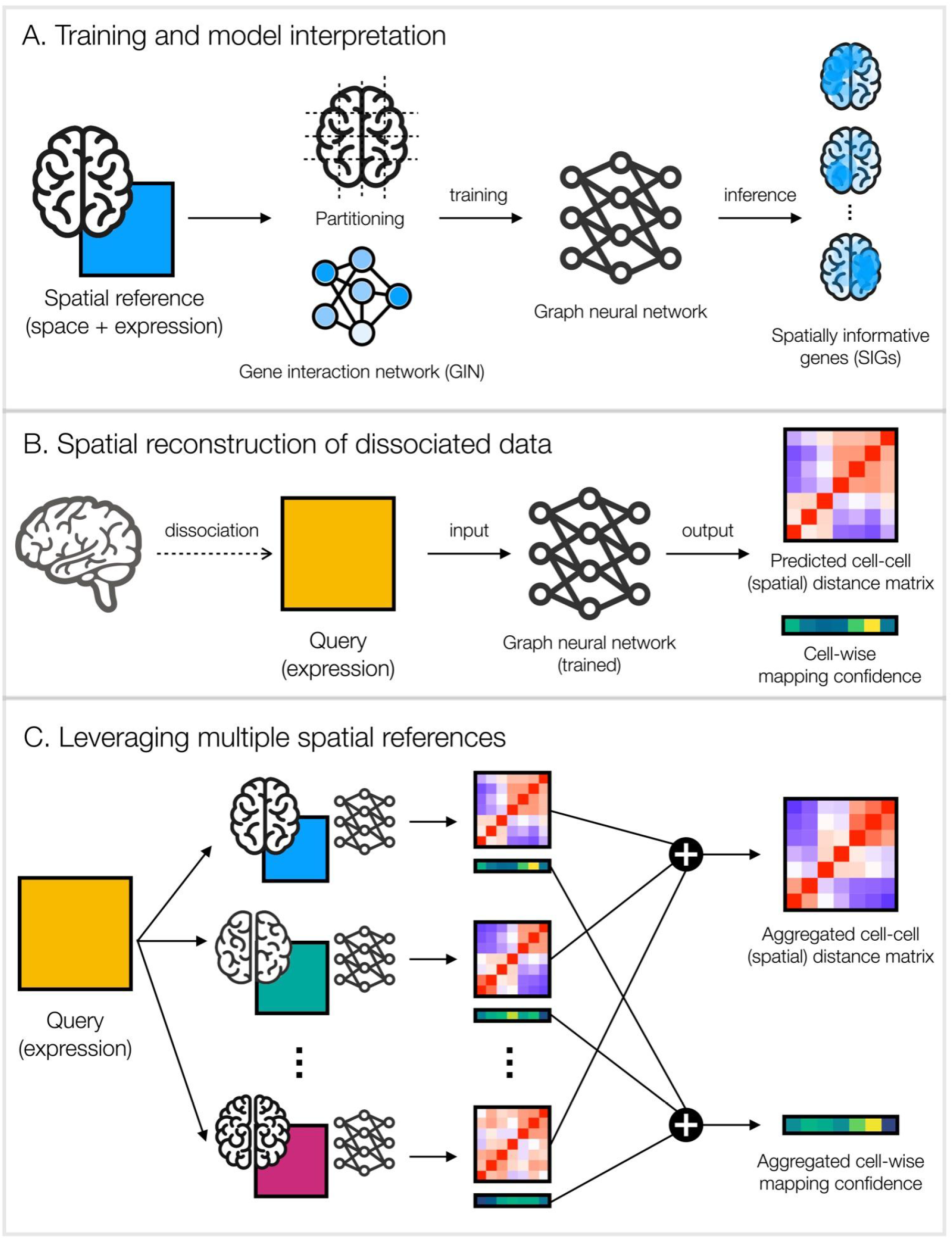
Mapping dissociated single cells onto their latent tissue of origin by classifying them to physical partitions of one or more spatially-resolved reference datasets. A. A spatial reference dataset, either cell-resolved or spot-resolved, is used to train the SageNet model. The spatial coordinates are first separated into non-overlapping partitions, and a gene interaction network is inferred from the gene expression data. These are used to train a graph neural network, which can be used to extract SIGs. B. Dissociated single-cell gene expression data are used as input into a trained SageNet model, for which a predicted cell-cell (spatial) distance matrix and a cell-wise mapping confidence vector is returned. C. SageNet models trained on multiple spatial references can be combined as an ensemble to generate aggregated cell-cell spatial distance matrix and cell-wise mapping confidence for a given query dataset.

Once the ensemble SageNet model is trained, a probability vector is estimated for each query cell, corresponding to each spatial partitioning (Methods). A spatial cell-to-cell distance matrix is calculated using the Jensen-Shannon divergence (Supplementary Figure 1B, (Endres and Schindelin 2003)), and is then used to embed the query cells in a 2 or 3-dimensional space; this low-dimensional space represents the SageNet spatial reconstruction (Supplementary Figure 1C). For each query cell, we additionally calculate a confidence score using Shannon’s entropy on the probability vectors (Figure 1, Supplementary Figure 1B-C), representing the degree of confidence in reconstructing the physical space. To facilitate interpretation of the SageNet reconstruction, we use global and local metrics including cell type co-location and mixing (Methods).

### SageNet recapitulates complex tissue patterning in early mouse organogenesis

To assess the relative performance of SageNet with other state-of-the-art approaches, we used a seqFISH dataset collected by (Lohoff *et al.* 2021) for cross-validation. This Spatial Mouse Atlas dataset contains barcoded gene expression measurements for 351 genes in three distinct mouse embryo sagittal sections (named embryos 1, 2, and 3), collected on embryonic day (E)8.5. Each section contains spatial gene expression for two distinct tissue layers 12 μm apart (denoted as layers 1 and 2). We assigned the cells from embryo 1 layer 1 as the spatial reference, and retained a random subset of cells from embryos 1, 2 and 3 to build a composite query dataset (Figure 2A, Methods). In this set-up, we were able to assess the quality of spatial reconstruction of SageNet in comparison to the true spatial coordinates of the query data. We found that SageNet, parametrised with five partitionings (Methods) and a GIN obtained using graphical LASSO (GLASSO) (Supplementary Figure 2), maintained a higher Spearman’s correlation of true and predicted cell-cell proximities (Figure 2B, Methods) across all embryos compared to other approaches including Tangram, novoSpaRc, and a naive approach using direct 2-dimensional embedding of the gene expression data (Methods). The improved quality of SageNet was especially clear for cells from the biologically independent embryos 2 and 3, suggestive of robustness to over-fitting and generalizability to new contexts. More broadly, we noted a more accurate reconstruction of the mixing relationships between cell types in the reconstructed space (Figures 2C,D, Supplementary Figures 3-4). Additionally, to investigate whether both the selection of genes and the graph neural network were contributing to the improved performance of SageNet, we used the set of SIGs selected by SageNet (Supplementary Figure 5) as input set for the naive PCA approach. This revealed that SageNet still showed an improved performance, albeit the improvement was smaller when using an improved set of input genes for the naive PCA approach (Supplementary Figure 6).

**Figure 2.**
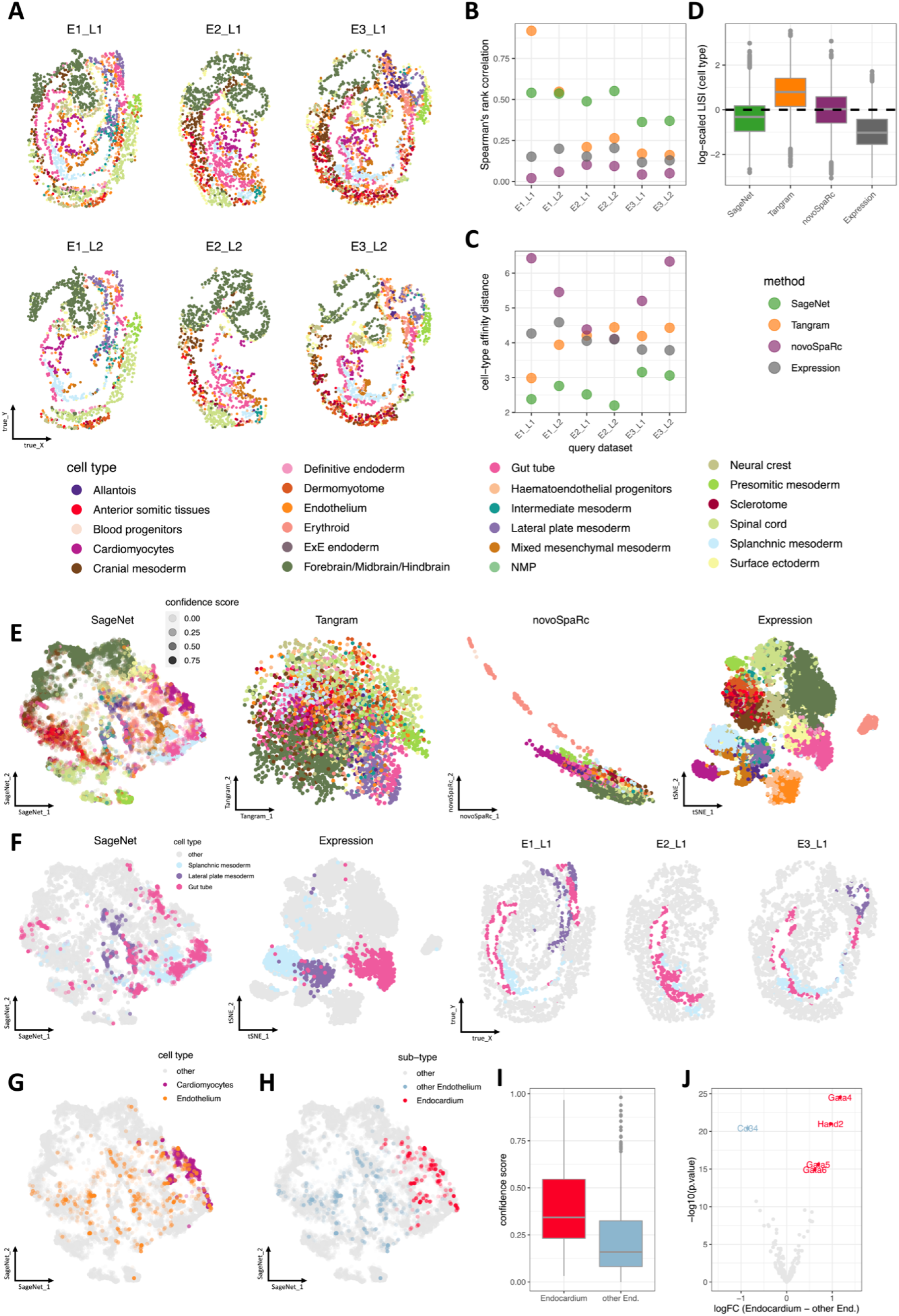
SageNet outperforms alternative reconstruction methods in cross-validation with the Spatial Mouse Gastrulation atlas A. Query cells selected from seqFISH-resolved embryo sections plotted in their true physical coordinates, coloured by cell types as in (Lohoff *et al.* 2021). B. Spearman’s rank correlation between each method’s predicted cell-cell distance matrix and the true spatial distance matrix, by embryo section. Higher values indicate a more accurate reconstruction. C. Matrix distances between each method’s predicted cell type contact map and the contact map computed using the true physical coordinates, by embryo section. Lower values indicate a more accurate reconstruction. D. Boxplot of log-scaled LISI ratios for cell type mixing; The LISI ratio for each cell is calculated by dividing the LISI score computed for the cell in each method’s reconstructed space by the LISI score computed using the true physical space. Values closer to 0 indicate a more accurate reconstruction. E. The reconstructed 2D space according to various methods; a subset of 10,170 query cells from the union of all 6 seqFISH-resolved embryos (20% of all cells from each embryo) is shown. Cells are coloured by cell type. Transparency of cells for SageNet are set according to their calculated mapping confidence level. F. SageNet recovers colocalization of gut tube and splanchnic mesoderm in mouse gastrulation. The reconstructed space according to SageNet is as in panel E and using gene expression only as in panel E, and the true physical space of embryos E1_L1, E2_L1, and E3_L1, ordered from left to right. G. SageNet reconstructed space as in panel E, subset to cardiomyocytes and endothelium cells. Transparency represents confidence of the mapping. H. SageNet reconstructed space as in panel E, subset to endothelium cells only. Cells are coloured by their sub-type annotation as Endocardium or Other Endothelium (Methods). I. Boxplots of SageNet’s confidence scores for the mapped endothelium cells, split by sub-type. J. Volcano plot showing the results of differential gene expression analysis on the sub-types of endothelium cells. The top five differentially expressed genes are highlighted.

Moreover, we visually noted SageNet’s ability to appropriately co-locate cells assigned different cell type labels in the reference in low-dimensional embeddings (Figure 2E). For example, gut tube and splanchnic mesoderm cells are transcriptionally distinct but proximal in the spatial reference; they are co-located in the SageNet embedding, but not in the other approaches (Figure 2F, Supplementary Figure 3).

Finally, since SageNet reports a mapping confidence score for each query cell, we were able to assess differences in mapping confidence within cell types (Supplementary Figure 7). Interestingly, we observed variation in mapping confidence among endothelial cells, with higher confidence for endothelial cells predicted to be closer to the developing heart tube (Figure 2G-I). The inner wall of the developing heart tube is lined with a specialised subset of endothelial cells, termed the endocardium (Dye and Lincoln 2020). The higher confidence of endothelial cells predicted closer to the heart, supports the hypothesis that these endothelial cells represent the endocardium, as evidenced by their distinct expression of genes such as *Gata4, Gata5, Gata6, Hand2,* compared to endothelial cells mapped to other regions of the embryo that exhibit a less distinct transcriptional profile (Figure 2J, (Dittrich *et al.* 2021; Duan *et al.* 2019)).

### SageNet accurately reconstructs the dissociated Mouse Gastrulation Atlas and recovers cell type colocalizations

Having demonstrated strong performance of SageNet, we next attempted to evaluate a spatial reconstruction of scRNA-seq data from the same stage of mouse development from which the spatial reference originated. To build the SageNet model, we first examined the suitability of an ensemble model incorporating all spatial references. Upon cross-comparison, we found that the ensemble SageNet model with embryos 1, 2, and 3 (Methods) was more robust across all embryos, with less over-fitting to any one spatial reference (Figure 3A).

**Figure 3.**
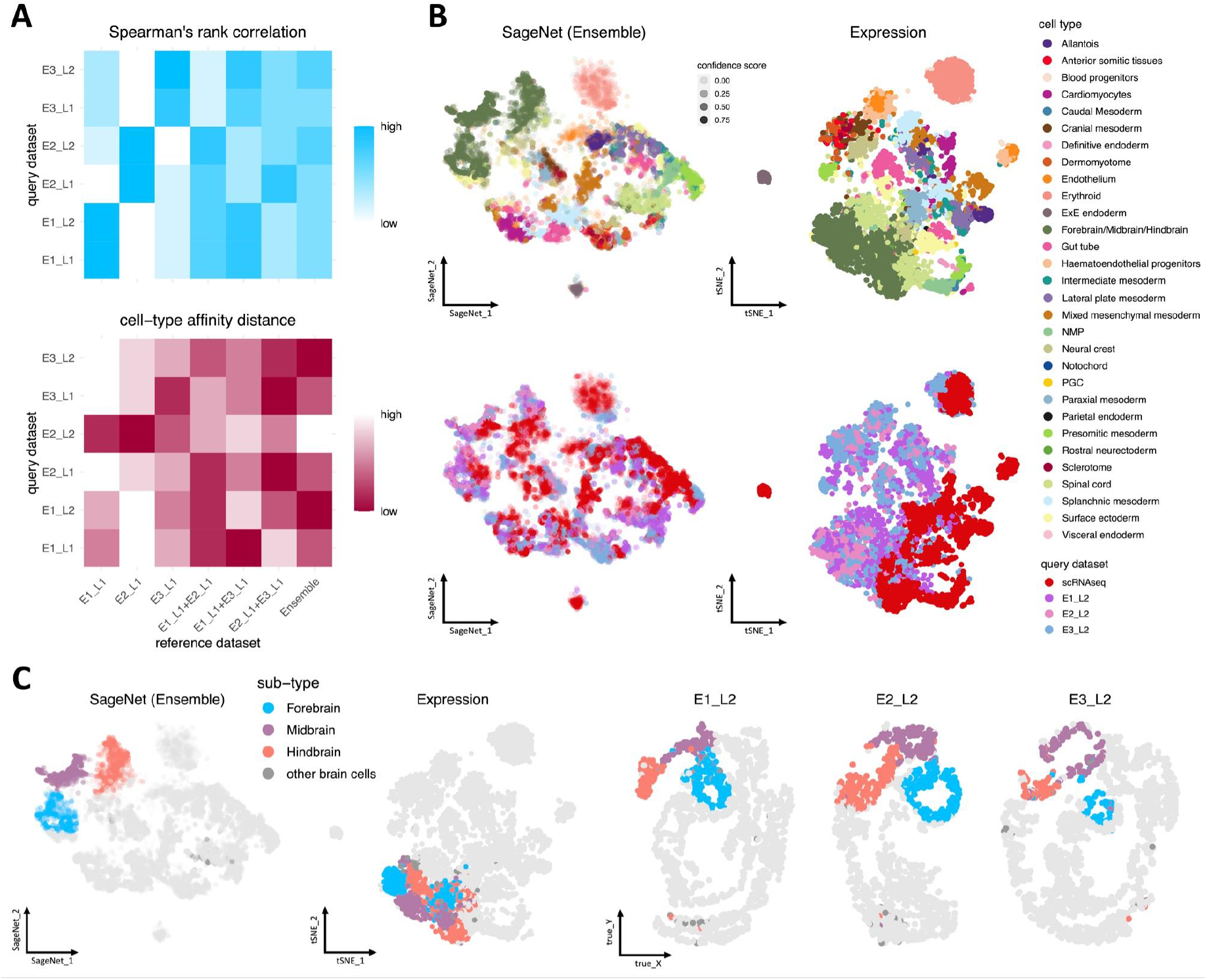
Superior performance of the aggregated SageNet model over individual models and spatial reconstruction of the dissociated Mouse Gastrulation Atlas A. Comparison of models trained on embryo 1 layer 1 (E1_L1), embryo 2 layer 1 (E2_L1), embryo 3 layer 1 (E3_L1) versus the aggregated SageNet models from pairs of embryo sections and all three embryo sections together (Ensemble). Each row represents a mapped query dataset, each column indicates a reference dataset combination, and colours represent performance, normalised by row. Top: Spearman’s rank correlation between the distance matrices computed by the SageNet models and the true spatial distance matrices. Bottom: the matrix distance between the cell type contact maps computed on the reconstructed spaces and the contact map computed on the true physical space. B. The reconstructed 2D space according to expression or aggregated SageNet model (Ensemble); a subset of 20% of cells is shown from the union of 3 seqFISH embryos, E1_L2, E2_L2, and E3_L2, and the scRNAseq mouse gastrulation atlas. Transparency of cells for SageNet reflects their calculated mapping confidence score. Top: coloured by cell type. Bottom: cells coloured by the corresponding query dataset. C. Brain cells are coloured based on hierarchical clustering on the reconstructed space by SageNet as shown in panel B. From left: SageNet’s reconstructed space; reconstructed space according to expression only; cells from the seqFISH query datasets shown in the true physical space.

We then proceeded to perform spatial reconstruction of a dissociated single-cell dataset using the better-performing ensemble SageNet model. Using a representative random subsample of approximately 4,200 cells from the dissociated scRNA-seq mouse gastrulation atlas, at embryonic day (E)8.5 (Pijuan-Sala *et al.* 2019), alongside a random subset of approximately 5,200 cells from the seqFISH layer 2, we generated a spatial reconstruction (Figure 3B). Interestingly, compared to a naive approach of expression-based reconstruction (Figure 3B), query cells were more mixed across samples according to data type (Supplementary Figure 8A-B), suggesting that SageNet is better able to account for platform-specific variation in reconstruction. We conclude that leveraging GINs as inductive bias to train GNNs makes SageNet robust and transferable to new unseen datasets (Embar, Srinivasan, and Getoor 2021).

By reclustering cells assigned as Forebrain/Midbrain/Hindbrain in the original study using the reconstructed SageNet space, we were able to determine distinct subgroups corresponding to the developing brain (Figure 3C). We found, consistent with existing knowledge (Martinez-Barbera *et al.* 2001), subregions of the developing brain were predicted to be near each other, specifically the forebrain nearer the midbrain, which in turn is near the hindbrain and tegmentum (Lohoff *et al.* 2021).

We noted a group of scRNA-seq derived blood/erythroid cells in the SageNet reconstructed space that exhibited a low confidence score (Figure 3B, Supplementary Figure 8C). The spatial reference data was depleted for circulating blood cells due to the tissue clearing step in the seqFISH experimental platform (Lohoff *et al.* 2021), thus it is worth noting that these blood-related cells are identified as having low mapping confidence, rather than the alternative of mistakenly being mapped to a different spatial region.

Taken together, these results demonstrate SageNet’s ability to build an accurate spatial reconstruction of dissociated single-cell data, revealing tissue organisation otherwise not represented using transcriptional information alone.

### SageNet builds a common coordinate framework using multiple spot-level spatial references and maps dissociated cells to their corresponding region in the developing human heart

To examine the performance of SageNet using spot-based spatial references (Longo *et al*.. 2021), we used a publicly available study of the developing human heart (Asp *et al..* 2019), where Spatial Transcriptomics (ST) data was collected for 9 heart sections, alongside dissociated scRNA-Seq data from the developing human heart at the same developmental stage. The ST heart dataset consists of a total of 3,115 spots (each containing approximately 30 cells) across nine tissue sections of one developing human heart at 6.5 weeks. We performed a leave-one-out cross-validation (LOOCV) scheme in which we leave one of the sections out as the query dataset and combine all other eight sections into one reference dataset. In order for different samples to have roughly the same anatomical orientation, we partitioned each section with a regular grid that is the smallest possible, covering all spots in the section. We performed 2×2, 3×3, and 4×4 square grids (Figure 4A, Supplementary Figure 9A). To balance the number of datapoints with the number of features, we subset the dataset to 500 highly variable genes (Methods). Recovering co-localization of regions illustrates accurate reconstruction of space for individual sections using our leave-one-out approach (Supplementary Figure 9A). Quantitative comparison with other methods revealed better performance of SageNet in terms of recovering cell-cell distances and local region mixing (Figure 4B-D). Similar to the mouse comparison, we observed that SIGs improved performance of the naive PCA approach (Supplementary Figure 9B-D).

**Figure 4.**
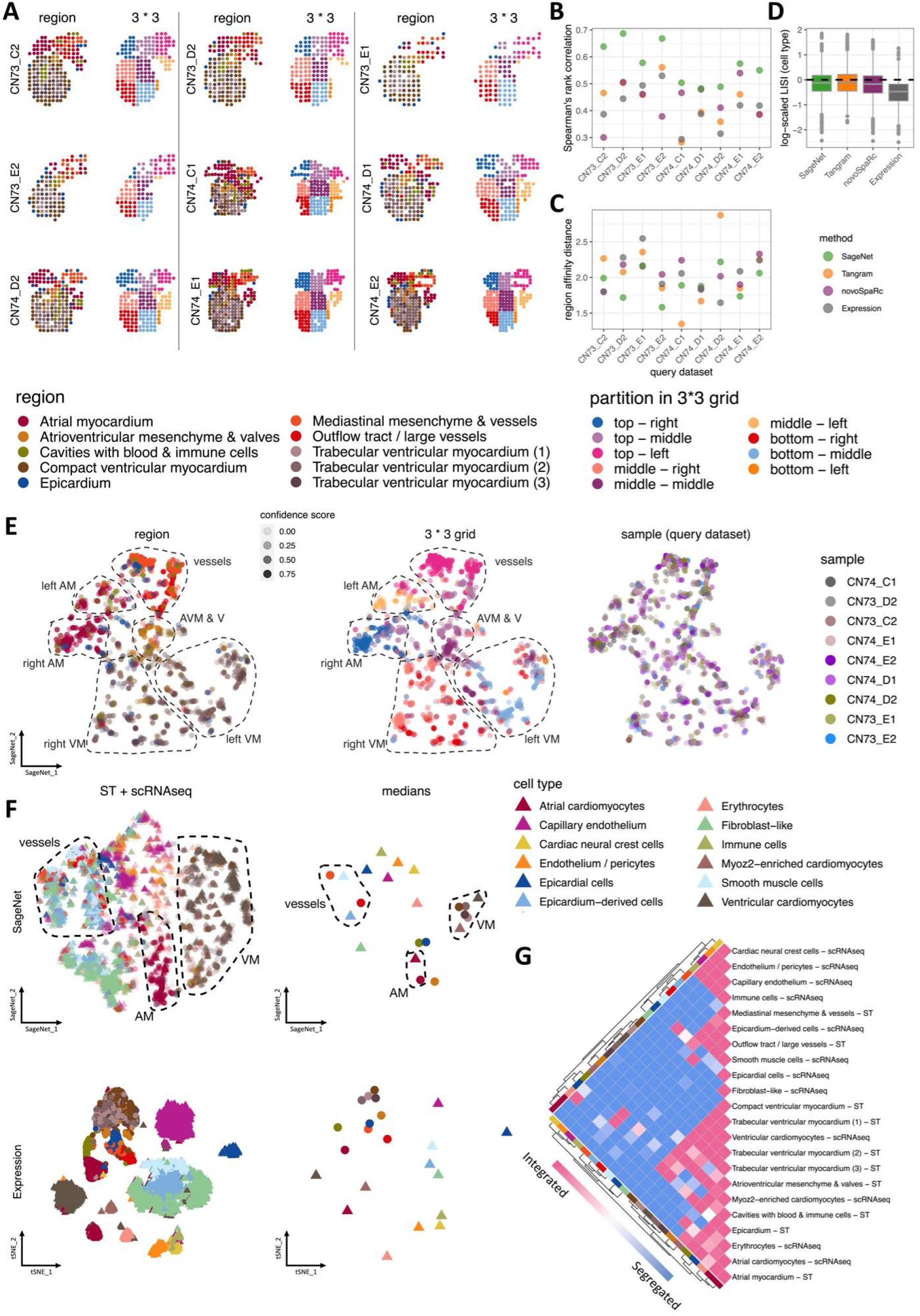
Sagenet robustly reconstructs the space and aggregates nonaligned spot-level Spatial Transcriptomics samples of developing human heart A. Spot-based Spatial Transcriptomics (ST) samples; each sample is shown with spots coloured by functional region given by (Asp *et al.* 2019) and the 3×3 grid partitions. B. Spearman’s rank correlation between each method’s predicted cell-cell distance matrix and the true spatial distance matrix, by sample. Higher values indicate a more accurate reconstruction. C. Matrix distances between each method’s predicted cell type contact map and the contact map computed using the true physical coordinates, by sample. Lower values indicate a more accurate reconstruction. D. Boxplot of log-scaled LISI ratios for cell type mixing; The LISI ratio for each cell is calculated by dividing the LISI score computed for the cell in each method’s reconstructed space by the LISI score computed using the true physical space. Values closer to 0 indicate a more accurate reconstruction. E. The joint 2D reconstructed space of the ST samples using SageNet; the model is trained on the whole spot-based ST dataset, with all samples being combined and the partitionings are as shown in panel A. Spots are coloured by left: region, middle: spot partitions from the 3*3 grids as shown in panel A, right: sample. Transparency of points is according to mapping confidence scores. F. Spatial integration of the ST and scRNAseq datasets of developing human heart; all cells and spots are embedded in the joint reconstructed space by SageNet (top row) and according to expression (bottom row). Right: median-based centroids of cell types and regions from the left column. Colours of solid circles correspond to regions as given in panel A, and colours of solid triangles correspond to cell types. G. Cell type/region contact map calculated on SageNet’s reconstructed space as shown in panel F. Cell types and regions are clustered based on the contact values by hierarchical clustering.

We used a trained SageNet model, on all 9 ST samples, to jointly embed the distinct ST samples into a common reconstructed space (Methods). Interestingly, we recovered the separation of functional subregions in the space such as the left and right atrial myocardium (AM), left and right ventricular myocardium (VM) and atrioventricular mesenchyme and valves (AVM & V) being mapped between atrial and ventricular myocardium, while also observing a high level of mixing of individual samples (Figure 4E). This indicates that SageNet is able to leverage multiple spatial references to build a common coordinate system for all tissue samples.

We used dissociated scRNA-seq data from the same developmental stage (Asp *et al.* 2019) to examine their latent spatial positions using SageNet. We found that the SageNet model, trained using spot-level ST data, was directly transferable to single-cell level based resolution with correspondence of known cell types including atrial and ventricular cardiomyocytes and their corresponding myocardium, and epicardium-derived and smooth muscle cells with vessels (Figure 4F-G). Remarkably, we observed a similar range of high mapping confidence for the unseen scRNA-seq cells as the reference ST-derived regions (Supplementary Figure 10). The epicardium is a multipotent population of cells known to give rise to different cardiac-related cell types during development including smooth muscle cells and fibroblasts. Interestingly, we found that the joint spatial reconstruction of scRNA-seq data and ST data resulted in the separation of epicardium-derived cells (EPDCs) into different regions, revealing further heterogeneity of EPDCs not identified in the scRNA-seq data alone (Figure 4F). One region of EPDCs most closely mapped to vessels and the outflow tract in the ST data and expressed markers of smooth muscle cells such as *OGN, ELN* and *CXCL12* (Supplementary Figure 11, (Cui *et al.* 2019)). Coincidentally, we found that the genes *ELN* and *CXCL12* were among the SIGs chosen by SageNet (Supplementary Figure 12). Conversely, another set of EPDCs most closely mapped with fibroblasts and were not observed to be closely aligned to a specific anatomical region, likely reflecting the presence of fibroblasts throughout the heart. Therefore, SageNet’s integration of ST and scRNAseq datasets unravelled three distinct regions for which epicardium-derived cells, fibroblasts, and smooth muscle cells were assigned. Taken together, these results suggest that SageNet is highly effective in integrating spatial and non-spatial data of different technologies and facilitates the understanding of developmental heart biology.

## DISCUSSION

We introduced SageNet, a computational approach that leverages spatially-associated gene-gene interactions to robustly reconstruct spatial relationships of dissociated single cell data, using gene interaction networks and graph neural networks. We observed superior performance of SageNet on both cross-validation and real-world tasks of spatial reconstruction on a mouse embryogenesis dataset. Our cross-validation and analysis of spot-based spatial reference data of the developing human heart demonstrates the ability of SageNet to perform data integration across a wide range of technologies.

SageNet incorporates uncertainty quantification and model interpretability. SageNet computes a mapping confidence score per cell based on the models’ outputs, which can be used to identify cells mapped with low confidence. Quantifying the degree of confidence for mapping is invaluable for downstream decision making (Abdar *et al.* 2021). A low confidence score might arise due to: (i) transcriptional exchangeability of cells in space (e.g., blood cells that are circulating); or, (ii) an incomplete spatial reference (e.g., extra-embryonic mesoderm cells in the mouse gastrulation data). To interpret trained SageNet models, SIGs are identified, which are the most informative genes in reconstructing the space. These SIGs can highlight important biological processes that govern spatial patterning of tissues, leading to further understanding, targeted panel design, and hypothesis generation.

One key choice when using SageNet is the initial spatial partitioning. This should be done such that proximal cells with different gene expression profiles are partitioned together, so that the graph neural network can determine the underlying gene expression signature that corresponds to their common pattern in space. Therefore, spatial partitioning should be performed using only the spatial coordinates, irrespective of the gene expression profiles. An additional key element of SageNet is the use of gene interaction networks (GINs). Our use of ensemble SageNet models facilitates the use of multiple distinct GINs for model building, meaning that multiple spatial references with different feature sets could be used jointly to map dissociated data, e.g., SageNet could be used to map dissociated CITE-seq data into a single reconstructed physical space informed by both an Imaging Mass Cytometry (IMC) resolved and ST-resolved reference. Currently, SageNet requires that the query data include all features in the underlying GIN, but it could be extended to require only a subset of features.

SageNet’s modular implementation enables users to aggregate multiple trained models on distinct spatial references, thereby improving spatial reconstructions without the need to perform a priori registration or alignment of the spatial references. As demonstrated in our analysis of the human heart dataset, this functionality facilitates efficient integration of multiple dissociated and spatially-resolved atlases, representing a viable approach towards building a common coordinate system across multiple spatially resolved datasets (Rood *et al.* 2019). In practical terms, this means that users can use a single existing model to predict new data and combine results without needing to refit the models. This is especially useful when query datasets become large. Nevertheless, SageNet’s pipeline has some computational limitations. For instance, in constructing the GIN using the GLASSO algorithm, runtime increases approximately quadratically with the number of genes, thereby necessitating an a priori selection of highly variable genes. Another limitation associated with both runtime and memory is computing the cell-cell distances for a query dataset containing many cells. Future development of efficient algorithms for computing Jensen-Shannon distances and storing sparse representations of these will overcome current runtime and memory limitations.

SageNet is a novel approach for spatial reconstruction that outperforms other state-of-the-art methods. Nevertheless, some challenges remain. For instance, it remains difficult to understand the fundamental limit for which spatial reconstruction methods can work, since such methods including SageNet assume that at least some spatial information is encoded in the transcriptional profiles of dissociated cells. Additionally, it is unclear how dissociated cells can be reliably mapped to spatial references with symmetric or repeated patterning, where cells are essentially interchangeable in space given their transcriptional profiles.

SageNet is provided as a user-friendly, well-documented, and open-source python package; we envision that SageNet will be applied to further query contexts such as pathological and cancer tissues, and spatial references generalised to other data modalities, such as spatial multi-omics (Liu *et al.* 2020), leading to discovery of novel biological patterns. We foresee the use of SageNet reconstructed embeddings as the basis for downstream analysis tasks including cellular dynamics (Lange *et al.* 2022) and inference of cell-cell interactions (Efremova *et al.* 2020).

## METHODS

### SageNet Workflow

SageNet probabilistically maps dissociated single cells to their latent tissue of origin by assigning the query cells to spatial partitions of one or more spatially-resolved reference datasets using an ensemble of graph neural networks. The workflow consists of the following steps.

#### Model building

##### Preprocessing the spatial reference(s)

SageNet requires at least one spatial reference linked to the dissociated data. The spatial reference data must contain user-supplied spatial partitionings. Such partitionings can be derived from the spatial coordinates using network clustering on the coordinates, e.g. Leiden clustering. Spatial partitionings are used by SageNet to perform probabilistic multi-class graph neural network classification towards reconstructing spatial coordinates. Here, a spatial partition is defined as a physically connected group of cells from the spatial reference, i.e., a connected subgraph of the k-Nearest Neighbours graph of cells in physical space. One or multiple spatial partitions can be provided by the user or derived from the spatial reference data.

Additionally, the spatial reference(s) should contain gene expression data, for which a gene interaction network is estimated. The gene expression data is assumed to be appropriately scaled and normalised (e.g., the logarithm of normalised counts). To build graph neural networks, SageNet requires either the user to supply an existing gene interaction network (GIN), or to input the spatial reference gene expression data. In the latter case, SageNet employs the graphical LASSO (GLASSO, (Friedman, Hastie, and Tibshirani 2008)), implemented in the function GraphicalLassoCV using the python package scikit-learn (Garreta and Moncecchi 2013), to estimate a sparse and modular gene interaction network. Default values of alpha = 0.5 and 0.75 (defining the grid for the regularisation parameter of the GLASSO algorithm) are used to build the GIN using internal cross-validation. The GIN is an unweighted and undirected network where nodes correspond to genes and edges correspond to presence of any type of interaction that may be a useful inductive bias for the classification of spatial partitions using SageNet.

##### Training the SageNet model

SageNet trains an ensemble classifier to estimate the spatial cell-to-cell distances and in turn map cells onto a scaleless 2D or 3D space. The ensemble classifier consists of several graph neural network classifiers, one for each partitioning of each spatial reference. A single classifier is trained by feeding in a gene expression matrix of size *c* × *g* as well as a gene-gene interaction network on *g* genes and outputs a *c* x *P* probabilistic classification, where *c* and *P* are the number of cells and partitions respectively.

###### Graph neural network classifiers

SageNet’s basic model leverages graph neural networks (GNNs) to perform discrete classification on dissociated cells. In this context, the gene expression profile from query cell *k* is represented as a graph *G_k_* = (*V_k_, E_k_*), with *g* nodes equal to the number of genes measured. Each node *i* from set of nodes *V_k_*, in *G_k_* has 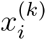 attribute representing one of *g* genes with a single scalar feature of gene expression *x*, and edges *E_k_* are unweighted and undirected with two nodes connected if the associated genes are connected in the gene interaction network (GIN). Thus, for all dissociated cells, the topologies of their graph representations are identical, but the attributes of nodes differ according to the cells’ gene expression profiles.

A graph neural network operation is defined as:

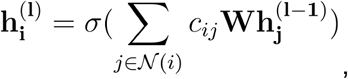

where 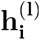 is the state of the node *i* after transformation in layer l, with the input values being 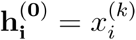 for *i* = 1,2,…, *g*. 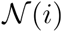 is the set of first order neighbours of node *i* in the graph, *c_ij_* is a learnable parameter that shows how much information should diffuse from neighbour *j* to node *i, σ* is a non-linear activation function, and finally *W* is another learnable parameter, which allows a global reweighting of information diffusion. By performing such operations, we simulate the information flow dynamics of gene interaction networks in shaping the morphology. The states of the final layer l = *L* are fed into a softmax layer for multi-class classification. The SageNet package provides implementation of multiple graph neural network classifiers via pytorch (Stevens, Antiga, and Viehmann 2020) and pytorch-geometric (Fey and Lenssen 2019).

In this study, we used ‘`TransformerConv`’ with one graph convolution layer of size 8 followed by one fully-connected layer of size 8. ‘`TransformerConv`’ uses graph transformer operator, *σ*, from (Shi *et al.* 2021) as:

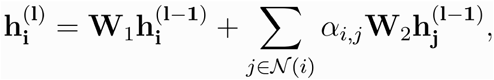

where 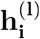 is the state of the node *i* after transformation in layer l, with the input values being 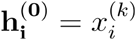 for *i* = 1, 2, …, *g*. The attention coefficients *α_i,j_* are computed via multi-head dot product attention:

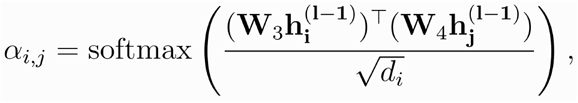

where **W**_1_, **W**_2_, **W**_3_, **W**_4_ are learnable parameters, 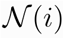 is the set of first order neighbours of node *i* in the graph, and *d_i_* is the degree of node *i*. Each model is trained for 20 epochs on a partitioning of a spatial reference.

##### Interpretation of trained model: finding spatially informative genes

We use the trained SageNet model to identify spatially informative genes. These genes can be interpreted as genes that are required for the accurate prediction of spatial partitions, and can be associated with specific partitions either in terms of presence of gene expression or importantly an absence of expression. To identify these genes, we first calculate the Integrated Gradients measure, *G_ijk_*, as implemented in captum (Kokhlikyan *et al.* 2020), for each gene *i*, each cell in the training data *j*, and each partition *k* across all partitionings. For each partition, the SageNet model determines a potentially nonlinear function *F*: ℝ^*n*^ > [0,1] which takes in a gene expression vector (of length n), for which a gradient for each gene 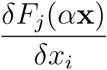 can be calculated, where *x_i_* is the ith element of the gene expression vector *x*. Thus we calculate the Integrated Gradients *G_ijk_* as

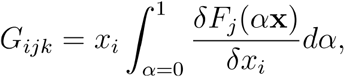

interpreted as the overall effect of perturbation of a gene’s expression to the resulting partition classification.

In turn, we aggregate across cells to estimate gene importance for each gene and partition, 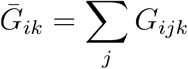, and subsequently extract the top *g* genes (by default 10) with largest magnitude across each partition, 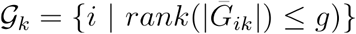. Finally, to determine the set of informative genes, we take the union of all such identified genes, 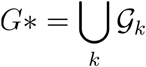. The number of informative genes may vary according to the choice of *g*, depending upon the degree of overlap of informative genes across multiple partitions and partitionings.

##### Combining, storing, and reusing trained SageNet models

Once a SageNet classifier is trained, the object can be saved and reloaded in a python environment. Since the SageNet model is an ensemble classifier, a loaded SageNet object can be subsequently appended to another SageNet object to extend the ensemble classifier. Other SageNet models may be trained using new spatial references, facilitating the inclusion of multiple spatial datasets for the task of spatial reconstruction. The spatial-omics technologies used to collect the spatial references could vary as long as the set of genes used in the reference dataset and the query dataset are the same. It is noteworthy that not all spatial reference datasets need to share the same set of genes, but the set of genes that each model (corresponding to a partitioning on a spatial reference) is trained on should be present in the query dataset.

SageNet models can be used to map new query data, allowing memory efficiency by only requiring the model object and query data, and not necessarily the training data, to be loaded in the environment. In addition, large query datasets may be split into multiple groups and inputted to SageNet, enabling the opportunity for parallelisation and to avoid memory limits.

### Mapping a query dataset

#### Preprocessing

The query dataset, typically a dissociated scRNA-seq dataset, consists of an appropriately scaled and normalised gene expression log-counts matrix with the same genes measured as in the spatial reference.

#### Probabilistic mapping

For each query cell input into each classifier of the ensemble SageNet model, the output is a numeric value between 0 and 1 for each partition. While the partition with the largest predicted value may represent the most likely partition according to the model, we aim to output a probabilistic mapping for all partitions. To do so, we perform the softmax transformation to estimate the relative probabilities associated with each partition, 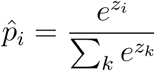, where *z_i_* is the model output for class *i*.

#### Quantifying mapping confidence

In addition to probabilistic mapping of query cells, we quantify the degree of confidence associated with the mapping of each query cell. Query cells may exhibit a low confidence score due to noisy gene expression, or mapping to multiple spatial partitions, and can be used to assess biological relevance. We calculate the confidence score of each query cell *k* using the Shannon’s entropy of the estimated probabilities for each model *j* within the ensemble SageNet model,

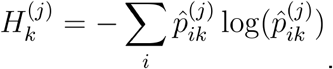

To estimate an overall confidence score for each query cell, we calculate the sum of entropies divided by their maximum possible value,

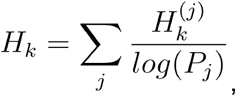

where *P_j_* is the number of partitions in the model *j*. Large values of *H_k_* indicate a low confidence for the mapping of cell *k*.

#### Cell-cell spatial distances

An advantage to probabilistic mapping of spatial partitions is the ability to calculate nuanced distances between cells, which can be used for further biological interpretation. We use the Jensen-Shannon divergence (JSD) (Endres and Schindelin 2003), a metric used to calculate the distance between two probability distributions, to calculate pairwise distances among query cells for each graph neural network of SageNet classifier *j*, 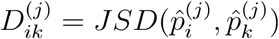, resulting in the distance matrix *D^k^*. We obtain a global cell-cell distance matrix *D* by calculating the element-wise sum of normalised distance matrices,

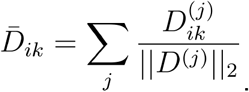

#### Low dimensional embedding

The global cell-cell distance matrix 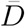 can be provided as input to non-linear low-dimensional embedding methods, such as t-SNE and UMAP, to obtain a 2D or 3D representation of cell-cell relationships. The low-dimensional embedding can be interpreted as a reconstruction of physical space since these approaches preserve short-range cell-cell distances. In this study, we used the t-SNE approach.

### Benchmarking SageNet

#### Benchmarked methods

We compared SageNet to four other approaches, described in more detail below. The same reference and query datasets were used for all methods.

##### Naive PCA approach according to only expression

Euclidean cell-cell distances are computed based on their gene expression values. Top ten principal components (PCs) were selected after performing a PCA on the log-normalised counts. The cell-cell distance matrix is then used by t-SNE implemented by R package Rtsne with perplexity 50 to embed cells in a 2D space. This approach represents a naive baseline, under the assumption that cells with similar gene expression profiles are likely to be located in the same regions in tissue space.

##### SageNet Markers (SageNet_SIG)

To understand whether SageNet facilitates understanding of genes that are more likely to be associated with spatial positioning, we identify the spatially informative genes according to the fitted SageNet model, using the default of 5 genes per partition per network. We then use these marker genes as input to the `Expression` approach as described.

##### novoSpaRc (novoSpaRc)

We use novoSpaRc (v0.4.3) under default settings to map query datasets to spatial references. novoSpaRc probabilistically maps each cell in the query dataset to all cells in the reference dataset according to optimal transport metrics. To evaluate and visualise, for each query cell we select the spatial coordinate of the reference cell with the highest estimated probability. Thus, for a query dataset we extract an embedding of the query cells in a 2D (or 3D) space.

##### Tangram (Tangram)

We use Tangram (v1.0.2) under default settings in “cells” mode to map query datasets to spatial references. Tangram trains a convolutional neural network to predict the best matching reference cell for a given query cell. The output is similar to novoSpaRc, that is, a probabilistic mapping of cells onto the reference cells/spots, from which we select the spatial coordinate of the reference cell with the highest estimated probability. The output is a 2D embedding of query cells.

#### Evaluation metrics

SageNet and the other methods used in our analysis provide a cell-cell distance matrix that is used to evaluate method performance at local and global scales. To compare method performance, we evaluate using the following criteria:

1. how accurate the true cell-to-cell distances are approximated (given that the ground truth exists);
2. how well cell type heterogeneity is preserved at local scale, i.e., around each cell; and,
3. to what extent cell type co-localization is recovered at the global level (given that the ground truth exists).

These evaluation metrics require ground truth, therefore we evaluate methods using spatially-resolved datasets as query datasets. We evaluate criterion 1 by comparing the predicted cell-cell distance matrix to the (true) physical cell-cell distance matrix (Euclidean distance). In cases where the cells in both reference and query datasets are labelled by cell types, we can evaluate criteria 2 and 3. For each evaluation criterion, we use the following metrics:

1. Spatial correlation We compute the Spearman’s rank correlation between lower-diagonal entries of the true physical cell-cell distance matrix and a methods’ cell-cell distance matrix. The rationale of using the rank-based Spearman’s rank correlation is to extract a robust measure of preservation of cell-cell proximities predicted by the methods.
2. Local Inverse Simpson’s Index (LISI) Given a cell annotation (e.g., cell type, batch, etc.) and embedding, LISI captures the degree of cell mixing in a local neighbourhood around a given cell. To evaluate how well local heterogeneity is preserved, we compute LISI as implemented in the R package Harmony (Korsunsky *et al.* 2019). When the ground truth is available, we compute LISI for cells in their true spatial locations as well. We then divide the LISI computed on the reconstructed space for each cell and divide it to the true LISI for that cell, finally we take the log2 of these values to compute the relative local heterogeneity scores.
3. Cell type and region contact maps and cell type affinity distance As introduced in (Lohoff *et al.* 2021), we compute cell type (or region) contact maps on the cell-cell (or spot-spot) distance matrices on the 2D reconstructed space, estimated by each method. To do so, we take the cell neighbourhood graph based on the cell-cell distance matrix and cell-level or spot-level annotation. We then randomly reassign the annotation by sampling without replacement (500 times) and calculate the number of edges for all pairs of annotated groups and compare it to the number of edges for which a particular pair of annotated groups was observed with the original annotation. To construct the cell–type (or region) contact map, we compare the randomly reassigned number of edges to the observed number of edges, across all permutations. Higher values correspond to the pair of annotation groups being more segregated, and lower values correspond to them being more integrated in reconstructed space than a random allocation. We then take a matrix distance between the estimated cell type (or region) contact maps and the true cell type contact maps from the reference dataset(s) and call this value cell type (or region) affinity distance. In case of multiple reference datasets, we take the element-wise mean to aggregate the true cell type contact maps.

### Comparison and analysis of molecule-resolved mouse gastrulation datasets

#### seqFISH study of mouse organogenesis

Lohoff *et al.* (2021) carried out a seqFISH experiment on sagittal sections from three mouse embryos corresponding to embryonic day (E)8.5–8.75 to quantify spatial gene expression at single cell resolution of a pre-selected set of 387 genes. For each embryo section, they captured two 2D planes, 12um apart, yielding a total of 6 spatially-resolved sections. The authors performed cell segmentation, quantified gene expression log-counts, and assigned cell type identities to each cell using a large-scale single cell study of mouse gastrulation (Pijuan-Sala *et al.* 2019) as a reference.

This dataset is ideal for evaluating SageNet in comparison with other methods, since we have access to ground truth spatial coordinates for individual cells over multiple biological replicates, and the tissue structure observed across mouse embryos are varied and complex. We downloaded the gene expression matrix and cell type and spatial location metadata from https://content.cruk.cam.ac.uk/jmlab/SpatialMouseAtlas2020/. Prior to analysis, we removed cells that were annotated as “Low quality” by the authors.

#### scRNA-seq study of mouse gastrulation and organogenesis

The mouse gastrulation atlas (Pijuan-Sala *et al.* 2019) is a 10x Genomics scRNA-seq dataset from mouse embryos spanning embryonic days E6.5-8.5. We used the Bioconductor package MouseGastrulationData (Griffiths and Lun 2021). For our analysis, we restricted the genes to those intersecting with the genes in the abovementioned seqFISH dataset, and removed cells that were annotated as “Doublet” and “Stripped”.

#### Composite query dataset

To understand the quality of spatial reconstruction and to facilitate biological interpretation, we built a composite query dataset of mouse organogenesis by selecting a random subset of 20% of cells (overall, 10170 cells) in each of all 6 seqFISH mouse embryos and 25% of the scRNAseq dataset (overall, 4227 cells), resulting in a total of 14,397 cells.

#### Cross-comparison of seqFISH samples

We took embryo 1 layer 1 as the spatial reference and partitioned with the leiden clustering algorithm (using squidpy) at 5 different resolutions, *r* = 0.01,0.05,0.1,0.5, and 1. We also estimated the GIN using the GLASSO, implemented by SageNet, with regularization parameters (0.5 and 0.75). Using this set of partitionings and GIN we trained an ensemble SageNet model consisting of 5 individual models. We then mapped the Composite query dataset using the aforementioned trained models and obtained predicted cell-cell distances. We also computed the Euclidean cell-cell distance matrix based on the 10 first PCs of the log-normalised gene expression values after performing PCA and also subset to the markers proposed by the SageNet model (only based on the reference dataset). We also obtained the 2D output of Tangram and novoSpaRc and computed the Euclidean cell-cell distance based on the 2D embedding. We then computed the evaluation metrics (distance correlation, cell type affinity distance, and LISI distribution) on the predicted cell-cell distance matrices and the 2D embeddings and the true physical distances and the real physical space split by each mouse embryo section.

#### Discriminating endocardium and other endothelial cells

To further investigate the difference between endothelium cells mapped closer to cardiomyocytes and other endothelial cells, we extracted the distance, as output from SageNet, of each endothelial cell to the nearest cardiomyocyte cell. We then assigned the nearest 10% of endothelial cells as putative endocardium and performed a differential gene expression analysis, using the ‘findMarkers’ function from R package scran (Lun, McCarthy, and Marioni 2016).

#### Investigating performance of the aggregated model in comparison to the single-reference models

We used the seqFISH samples embryo 1 layer 1 (E1_L1), embryo 2 layer 1 (E2_L1), and embryo 3 layer 1 (E3_L1) as spatial references. We used the GLASSO approach to estimate the gene interaction network. To extract multiple spatial partitionings for each seqFISH sample, we obtained a spatial kNN graph using the implementation in squidpy, and performed Leiden clustering on the graph with resolutions *r* = 0.01, 0.05, 0.1, 0.5, and 1. For each seqFISH sample and partitioning, we fitted a SageNet model. We appended SageNet models across various partionings to generate ensemble SageNet models for each seqFISH sample, which we named according to the sample. We also appended all models to generate an ensemble SageNet model (‘`Ensemble`’). We then mapped the Composite query dataset, subset to seqFISH datasets, using each of these 4 models, and then compared them with respect to distance matrix correlations, cell type affinity distances.

#### Spatial reconstruction of scRNA-seq derived mouse gastrulation cells

We subset the composite query dataset to cells from the scRNAseq data and embryos 1 layer 2, 2 layer 2, and 3 layer 2 (as the spatial anchors) to be used as the query dataset for this task. We used the ‘`Ensemble`’ SageNet model described above to map this query dataset. We then computed the reconstructed space and cell type contact map on this reconstructed space.

#### LISI scores for batch and cell type mixing

We computed the log2 LISI scores based on the batch (seqFISH vs. scRNAseq) and cell type labels, in order to assess the level of cell mixing in the reconstructed space by the naive low-dimensional embedding approach and SageNet.

#### Spatial decomposition of brain

In the SageNet reconstructed space using the ‘`Ensemble`’ model we captured 3 clusters of brain cells. We performed hierarchical clustering according to Euclidean distances in the reconstructed space using ‘`hclust`’ with default parameters in R. We used cutree with 10 groups and then took the largest 3 clusters. We visually inspected the spatial positions of the cells belonging to these 3 clusters, and noted their distinct anatomical organisation, corresponding to the Forebrain, Midbrain and Hindbrain.

### Comparison and analysis of spot-level developing human heart datasets

#### Spatial Transcriptomics (ST) study on developing human heart

Asp *et al.* (2019), performed a study of developing human heart collected using ST along the dorsal-ventral axis from an embryo collected at 6.5 post-conceptional weeks. ST is a spot-level spatial technology, where each observation typically contains aggregated expression information from a small number of cells.

The ST heart dataset consists of a total of 3,115 spots (each containing approximately ~30 cells) across nine tissue sections of one developing human heart at the 6.5 week embryonic stage. We downloaded the normalised gene expression data along with region labels and spatial coordinates from the publication website (https://www.spatialresearch.org/resources-published-datasets/doi-10-1016-j-cell-2019-11-025/). Prior to analysis, we restricted the genes to the top 500 highly variable genes as obtained by the Bioconductor package scran (Lun, McCarthy, and Marioni 2016).

We generated consistent partitionings for each ST sample by assigning spots to positions in a square grid. Since ST spots are generated on approximately regular, evenly-spaced positions, we assigned spots to each partition according to their vertical and horizontal position in a 2×2, 3×3, and 4×4 square grid, resulting in three distinct partitionings. We used the GLASSO to estimate a global GIN on the whole ST dataset.

#### scRNA-seq study of developing human heart

Alongside the ST dataset, Asp *et al.* (2019) performed scRNA-seq on a set of 3,717 developing human heart cells from a biological replicate of the 6.5 ST sample. We downloaded the normalised gene expression log-counts and cell type labels for this data, and restricted it to the same set of genes for which we built the SageNet models.

#### Cross-comparison of ST heart samples and the leave-one-out schema

Due to the low number of data points, we were not able to take each ST section as a separate reference. Therefore, we performed a leave-one-out (LOO) scenario, where we left one of the sections as the query dataset and combined all 8 other sections as the reference dataset. The same partitionings as described above were used to train the model. We trained one ensemble SageNet model (for 3 different partitionings) on each of the sets of 8 reference datasets and mapped the corresponding query dataset. We followed the same procedure for the other methods, i.e., expression, SageNet SIG, Tangram, and novoSpaRc, and computed evaluation metrics as detailed above.

#### Spatial reconstruction of the ST dataset, scRNAseq dataset, and integration of ST and scRNAseq datasets

We trained the SageNet model using the entire ST dataset as the spatial reference with the abovementioned grid partitionings and a GIN estimated on the whole gene expression matrix across different samples (Supplementary Figure 12). We then reconstructed all ST samples jointly in a 2D space using this model. We also mapped the dissociated scRNA-seq dataset of developing human heart, from the same developmental stage, using this trained model as described above. Finally, we mapped the composite dataset, including both ST and scRNAseq cells, onto two dimensions using this model.

## DATA AVAILABILITY

All data files, including raw datasets, intermediate objects produced by SageNet (cell and gene metadata), benchmarking experiments’ outputs, pre-trained models, and final ggplot figure objects can be found at https://zenodo.org/record/6400952 (DOI: 10.5281/zenodo.6400951).

## CODE AVAILABILITY

We perform evaluation in R using the output files from python scripts implementing SageNet and the other benchmarked methods. SageNet is provided as an open-source python package. The package documentation including use cases and tutorials can be found at https://SageNet.readthedocs.io and the source package is available on github at https://github.com/MarioniLab/SageNet (DOI: 10.5281/zenodo.6394407). The scripts and notebooks for benchmarking experiments can be found at https://github.com/MarioniLab/SageNet_paper (DOI: 10.5281/zenodo.6402296).

## ACKNOWLEDGEMENTS

We thank our colleagues in University of Zurich, University of Cambridge Cancer Research UK Cambridge Institute, and the EMBL European Bioinformatics Institute for their support and intellectual engagement. We thank N. Yayon, E. Dann, A. Missarova, K. Hua, A. Richard, and other members of the Marioni lab for discussions concerning the project.

## AUTHOR CONTRIBUTIONS

E.H., S.G., M.R., and J.C.M. conceived the study. E.H. developed the method and software and performed data analysis with supervision from S.G., M.R., and J.C.M.. E.H. generated figures with supervision from S.G., M.R., and J.C.M.. E.H. interpreted the results with input from S.G., T.L., R.C.V.T, and J.C.M.. E.H., S.G., J.C.M. and M.R. wrote the manuscript. All authors read and approved the final version of the manuscript.

## FUNDING

The following sources of funding are gratefully acknowledged. E.H. was supported by EMBL-EBI (core funding to J.C.M.). M.D.R. and E.H. acknowledge funding from a collaborative project with Hoffmann-La Roche. T.L. was funded by the Wellcome Trust 4-Year PhD Programme in Stem Cell Biology and Medicine and the University of Cambridge, UK (203813/Z/16/A and 203813/Z/16/Z). R.T. was supported by the British Heart Foundation Immediate Fellowship (FS/18/24/33424). S.G. was supported by a Royal Society Newton International Fellowship (NIF\R1\181950) and by a Wellcome Trust Collaborative Award (220379/B/20/Z to J.C.M.). J.C.M. acknowledges core funding from EMBL and core support from Cancer Research UK (C9545/A29580). M.D.R. acknowledges support from the University Research Priority Program Evolution in Action at the University of Zurich and the Swiss National Science Foundation (grant number 310030_175841).

The funding sources mentioned above had no role in the study design; in the collection, analysis, and interpretation of data, in the writing of the manuscript, and in the decision to submit the manuscript for publication.

This research was funded in whole, or in part, by the Wellcome Trust. For the purpose of Open Access, the author has applied a CC BY public copyright licence to any Author Accepted Manuscript version arising from this submission.

## COMPETING INTERESTS

The authors declare no competing interests.

---

**Supplementary Figure 1.**
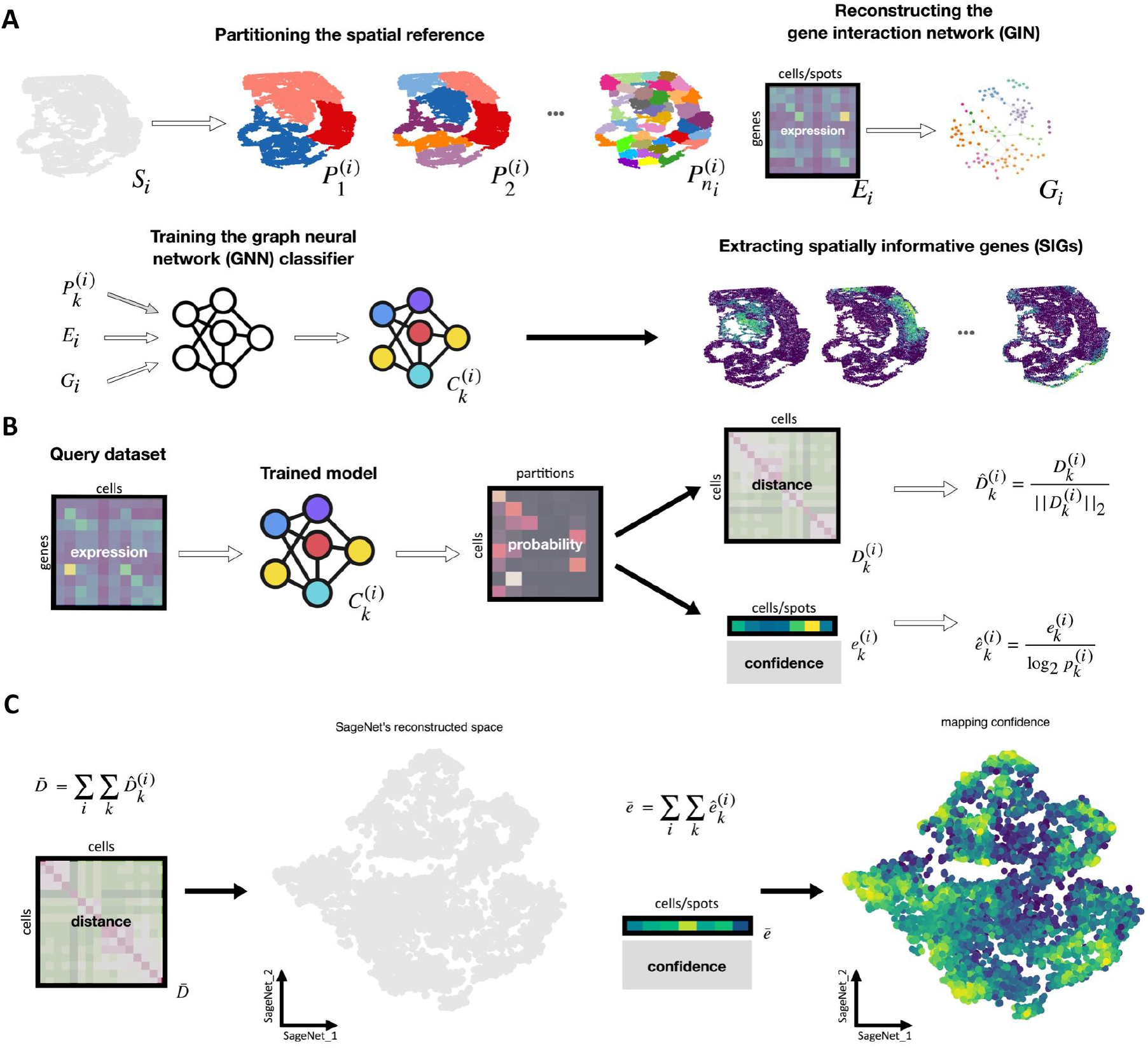
Mapping dissociated single cells onto their latent tissue of origin by classifying them to physical partitions of one or more spatially-resolved reference datasets. A. Training the models for the spatially-resolved reference dataset *S_i_*; *S_i_* is partitioned into physically disjoint regions based on the cells’ (or spots’) coordinates. The partitioning, 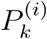, is performed *n_i_* times at different resolutions. The gene interaction network, *G_i_*, is constructed based on the gene expression matrix, *E_i_*. The graph neural network, 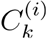, is trained to classify cells to partitions 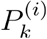, using *E_i_* as the input features and *G_i_* as the input graph structure. After training, the most important features are extracted from the trained model for each partition 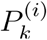. B. A query dataset is given to the trained classifier, 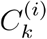. The classifier outputs a probability vector of size 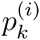 for each cell in the query dataset, where 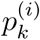 is the number of partitions in 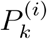. Pairwise distances between cells are computed using the Jensen-Shannon distance, giving 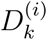. By computing the entropy of the probability vectors, the level of mapping confidence for each cell is computed as 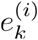. 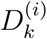 and 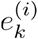 are normalised to 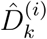 and 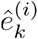 to ensure consistency among partitions of the other reference datasets. C. The normalised distance matrices computed on the outputs of all classifiers are added up to compute a global distance matrix 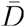. The normalised confidence scores are added up to compute a global mapping confidence, 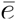 for the query cells. Using the computed distance matrix, one can compute a 2- or 3-dimensional embedding of the query cells.

**Supplementary Figure 2.**
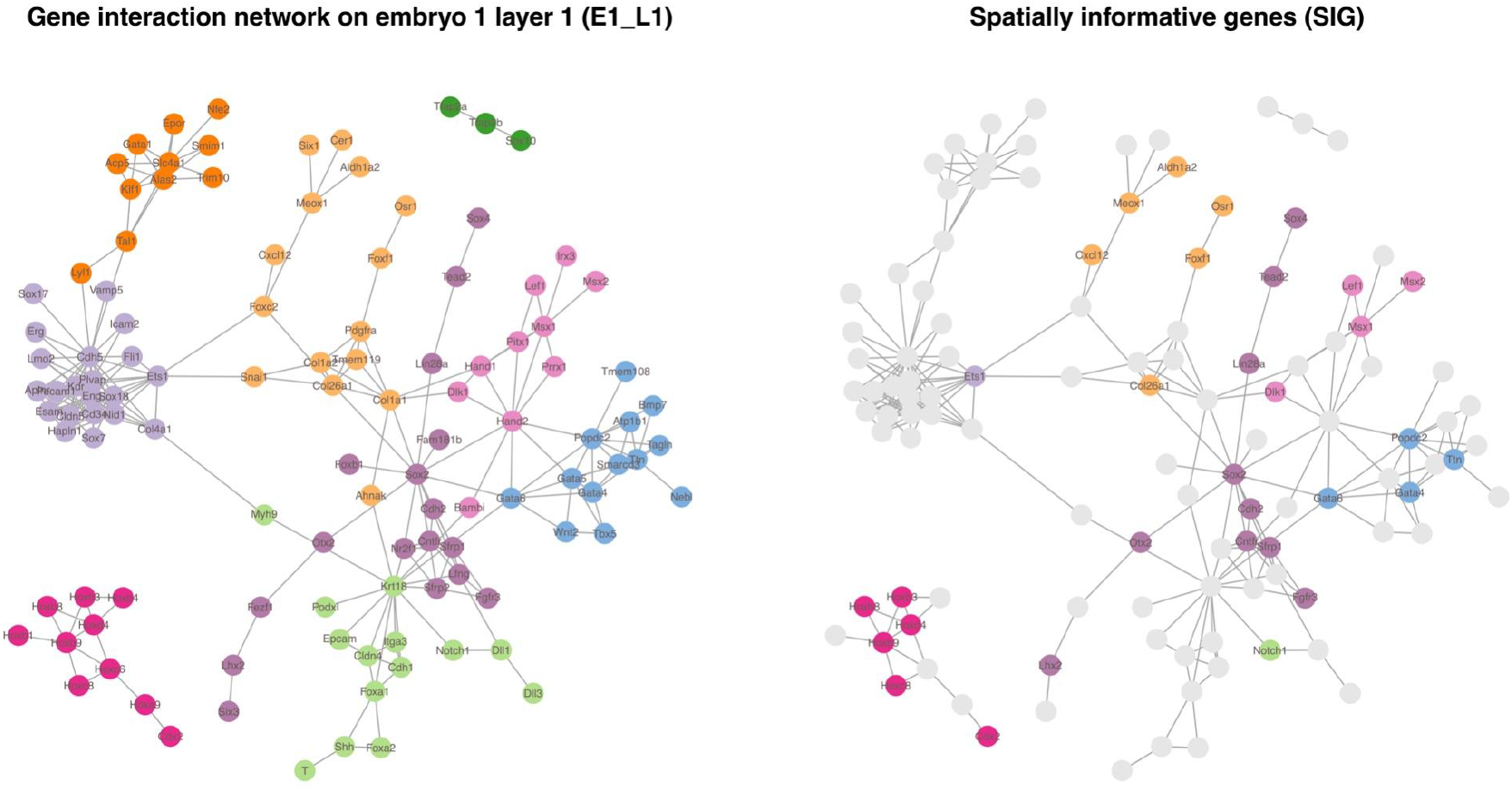
The gene interaction network on seqFISH reference dataset (Figure 2A), embryo 1 layer 1 (E1_L1). The gene interaction network (GIN) built on the reference dataset corresponding to Figure 2 (i.e., E1_L1); Each node represents a gene and each edge represents the statistical relevance (positive partial correlation estimated by GLASSO) of the two genes. Left: Isolated genes are removed and other genes are coloured based on their gene modules according to Leiden clustering. Right: SIGs inferred by SageNet are highlighted in the graph.

**Supplementary Figure 3.**
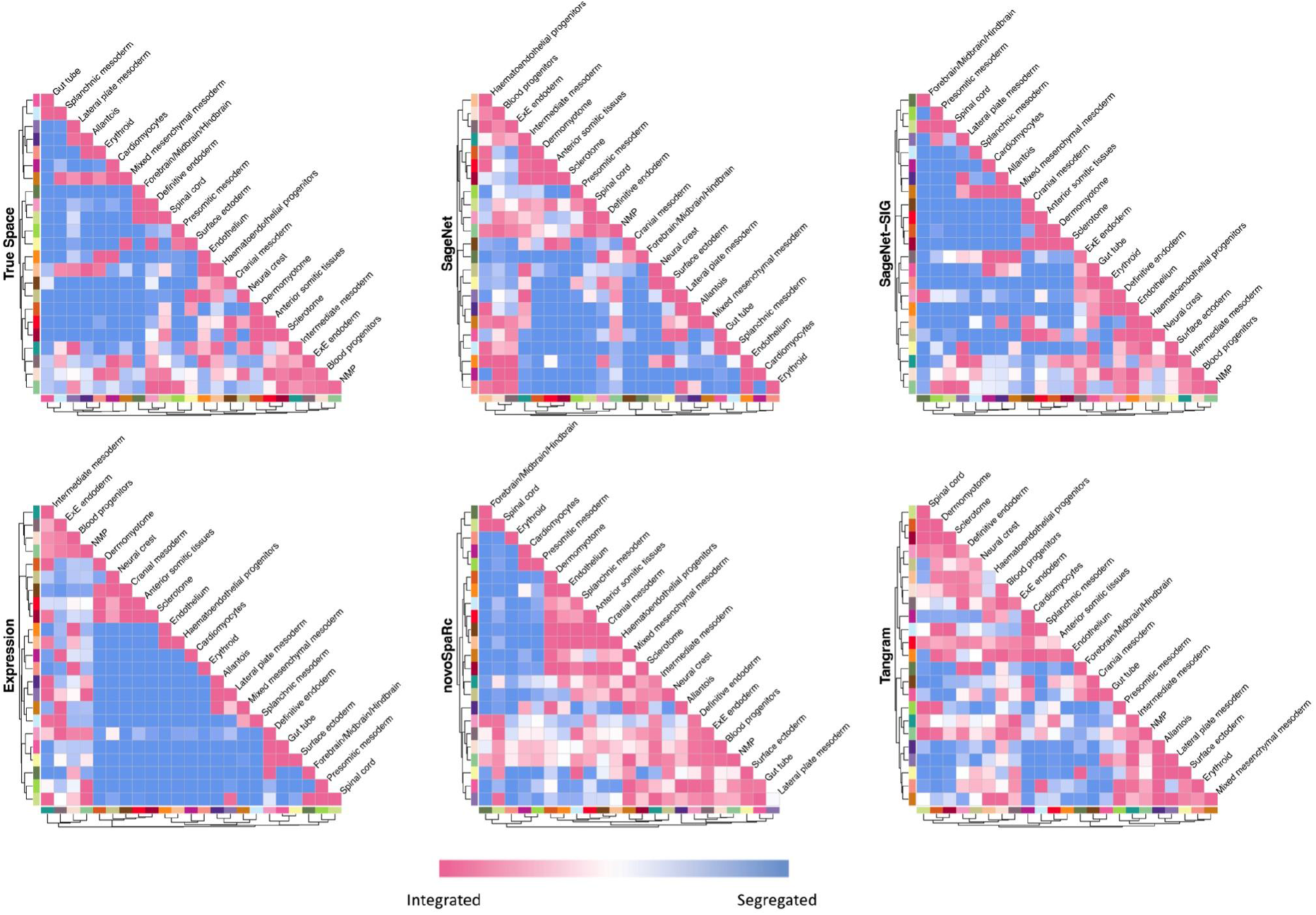
Cell type contact maps on the reconstructed spaces (shown in Figure 2E) using different methods for mouse embryo data. Cell type contact maps computed on the true physical space, and on reconstructed spaces using various methods. Cell types are clustered based on the contact values by hierarchical clustering.

**Supplementary Figure 4.**
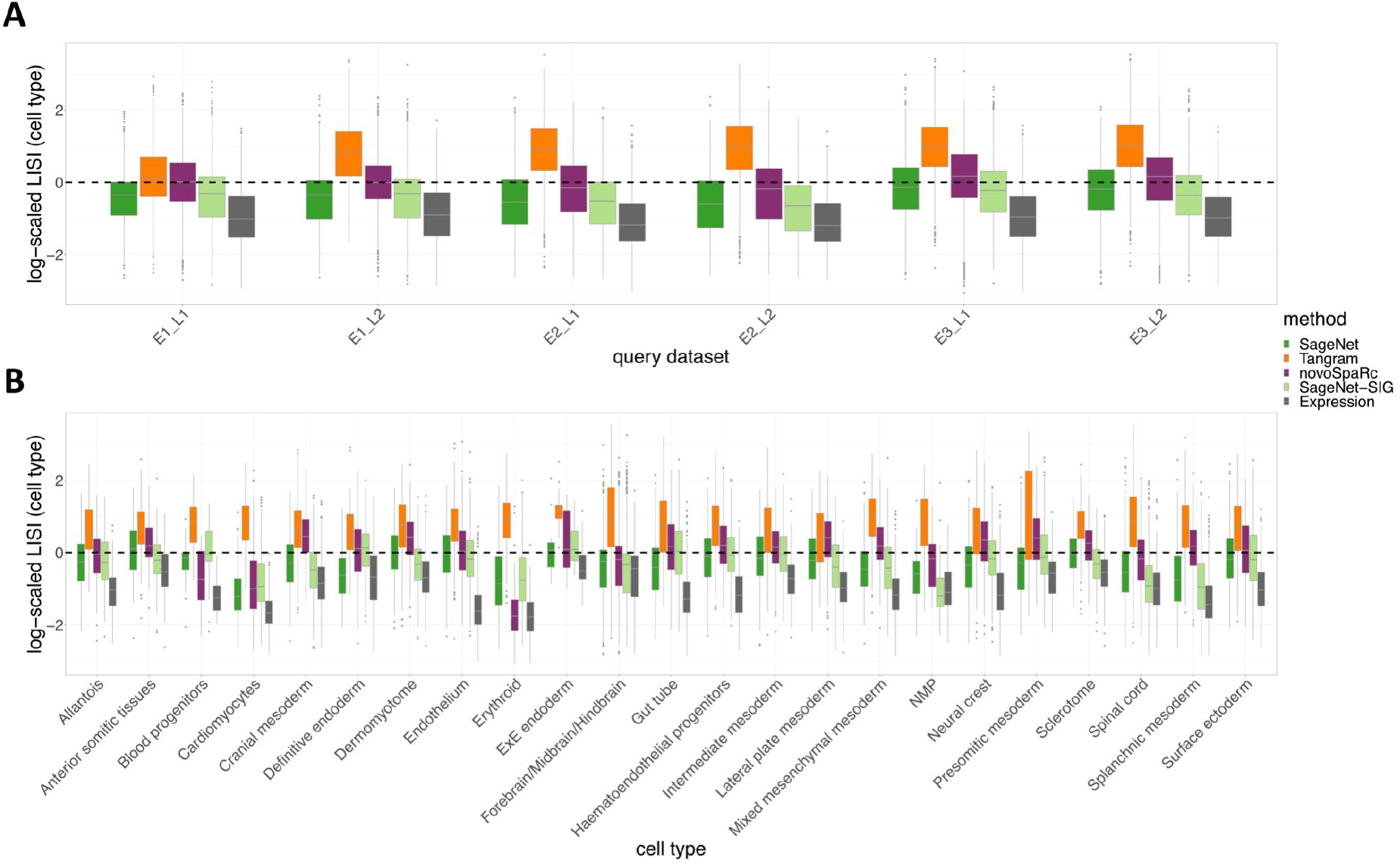
Scaled LISI scores for cell type mixing for mouse embryo data. A. Boxplot of log-scaled LISI ratios for cell type mixing; The LISI ratio for each cell is calculated by dividing the LISI score computed for the cell in each method’s reconstructed space by the LISI score computed using the true physical space. Values closer to 0 indicate a more accurate reconstruction. Boxplots are split by query dataset. B. As described in panel A, with boxplots split by cell type.

**Supplementary Figure 5.**
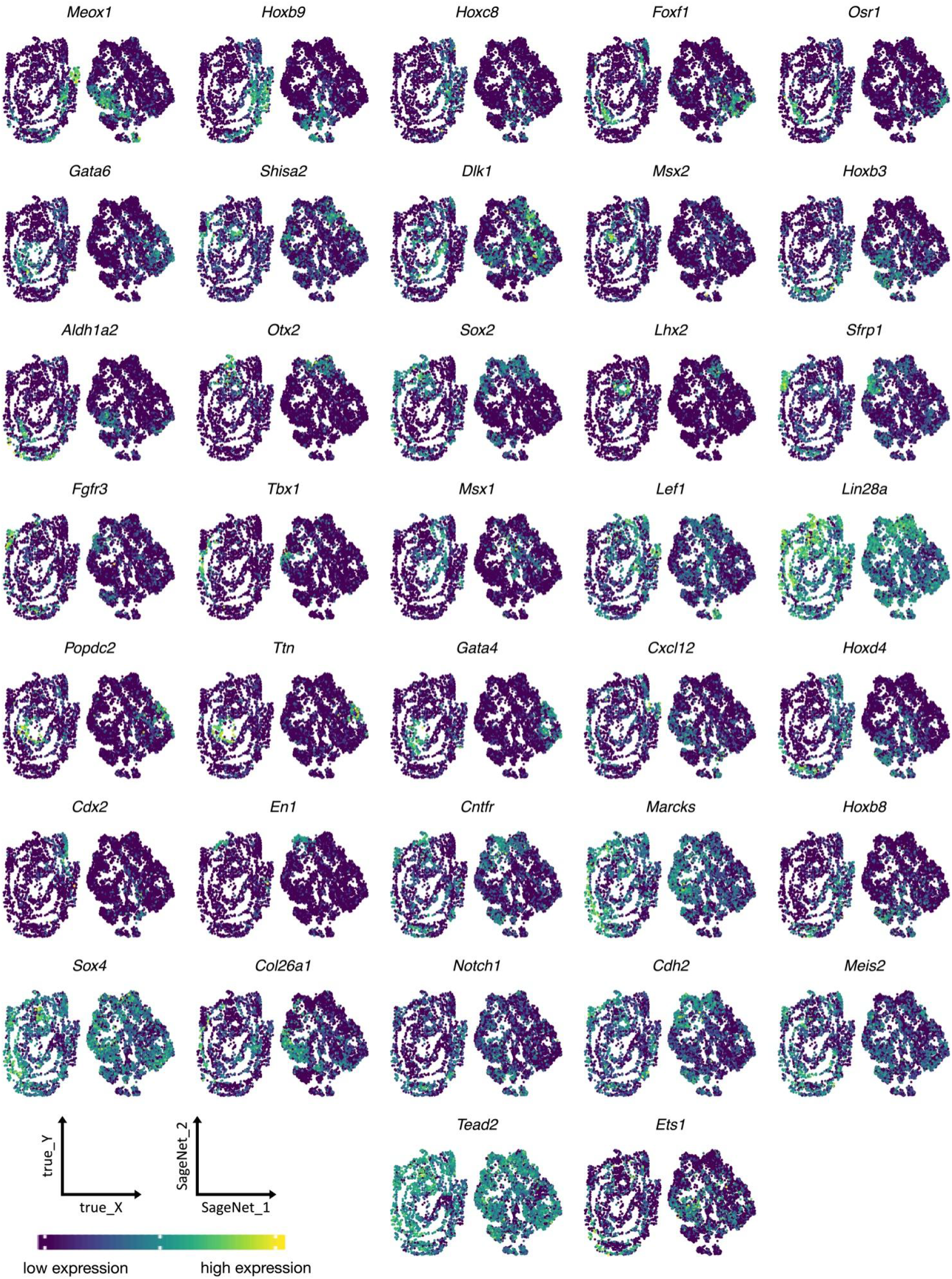
Spatially informative genes for mouse embryo data. Each panel displays gene expression of an SIG found by SageNet; left panels: embryo 1 layer 1 (E1_L1), right panels: the SageNet reconstructed space as shown in Figure 2E.

**Supplementary Figure 6.**
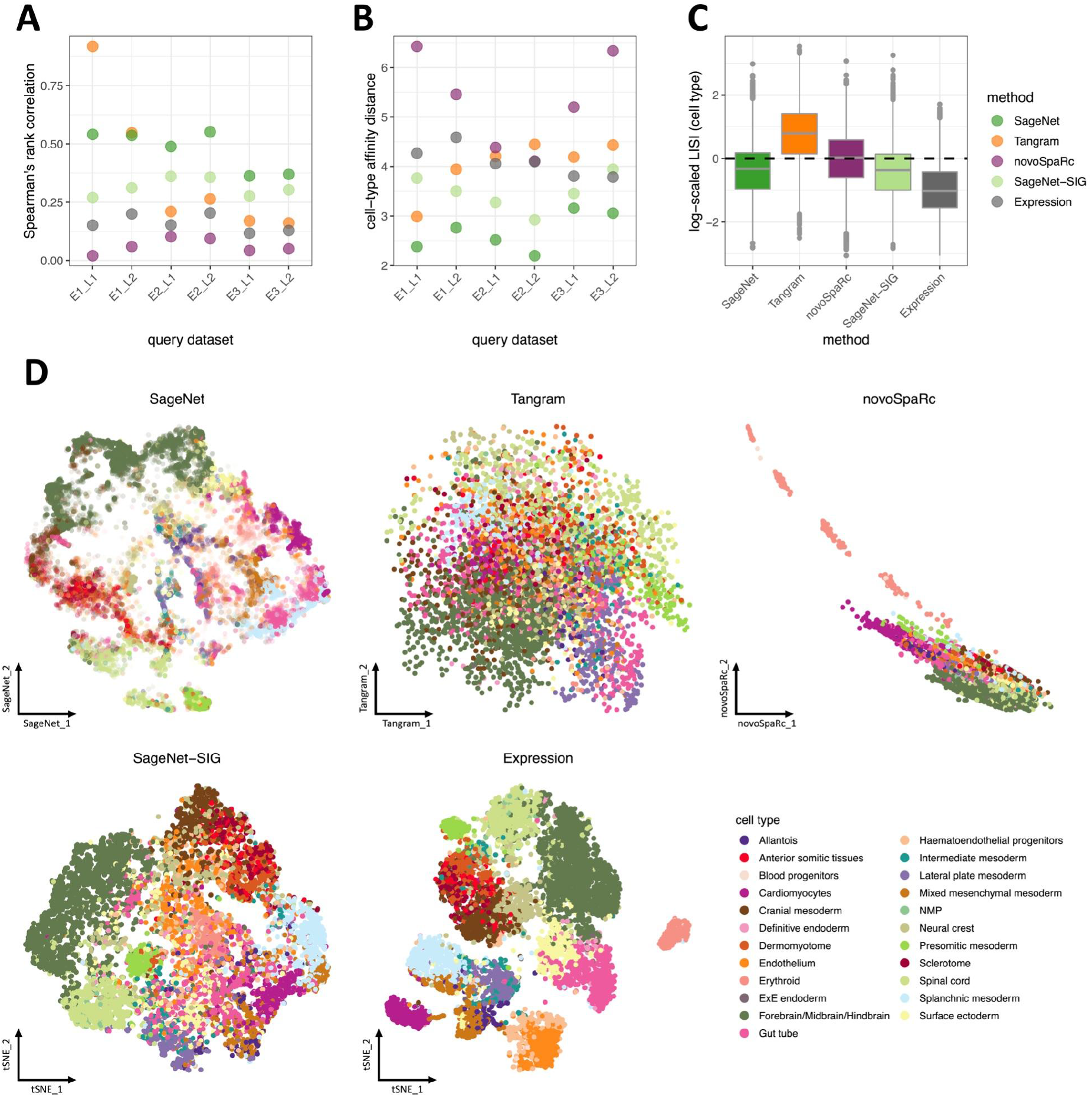
SageNet and SageNet-SIG outperform alternative reconstruction methods in cross-validation with the Spatial Mouse Gastrulation atlas A. Spearman’s rank correlation between each method’s predicted cell-cell distance matrix and the true spatial distance matrix, by embryo section. Higher values indicate a more accurate reconstruction. B. Matrix distances between each method’s predicted cell type contact map and the contact map computed using the true physical coordinates, by embryo section. Lower values indicate a more accurate reconstruction. C. Boxplot of log-scaled LISI ratios for cell type mixing; The LISI ratio for each cell is calculated by dividing the LISI score computed for the cell in each method’s reconstructed space by the LISI score computed using the true physical space. Values closer to 0 indicate a more accurate reconstruction. D. The reconstructed 2D space according to various methods, including SageNet-SIG; a subset of 10,170 query cells from the union of all 6 seqFISH-resolved embryos (20% of all cells from each embryo) is shown. Cells are coloured by cell type. Transparency of cells for SageNet are set according to their calculated mapping confidence level.

**Supplementary Figure 7.**
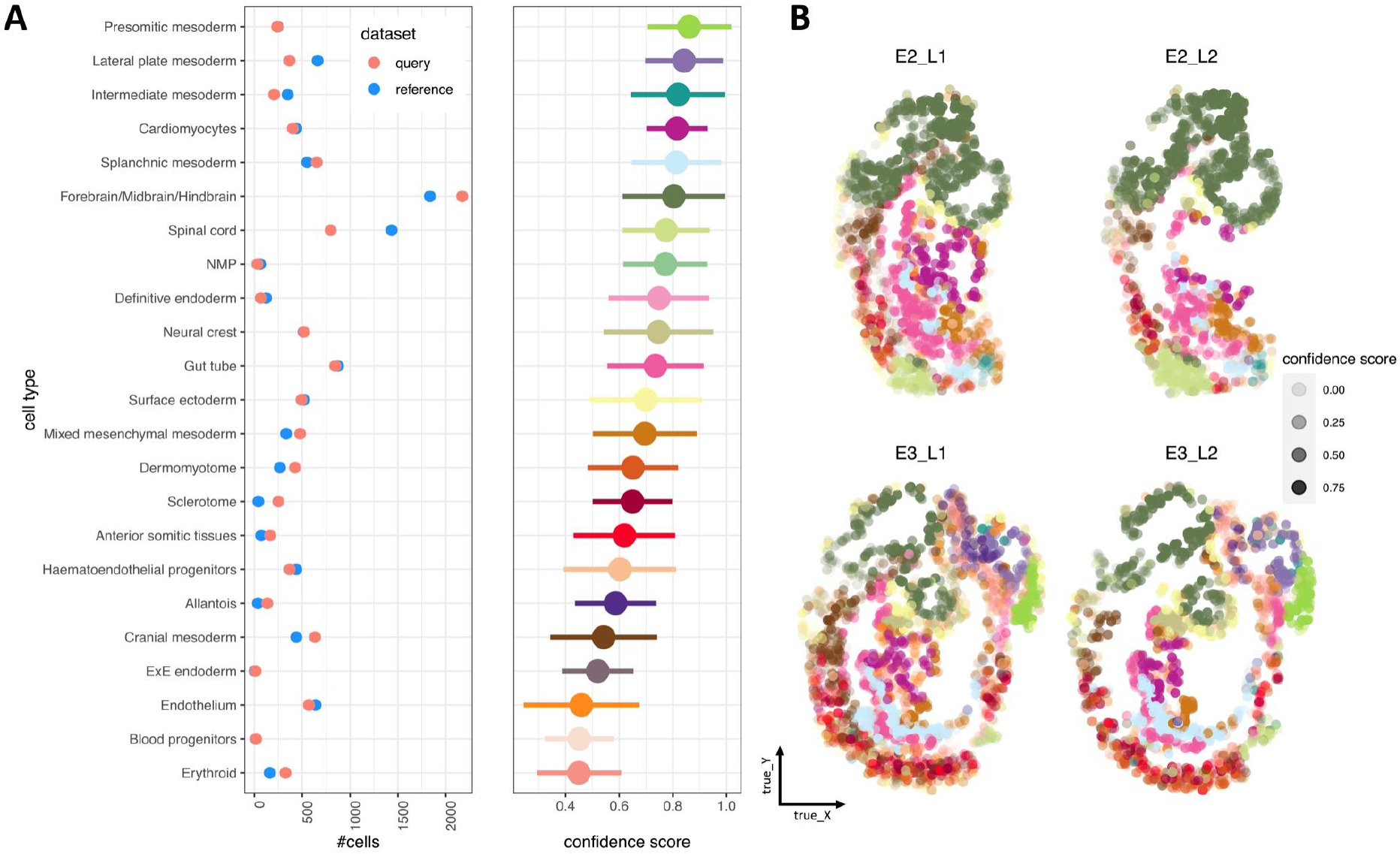
Mapping confidence scores of the SageNet model trained on embryo 1 layer 1 (E1_L1). A. Distribution of SageNet’s mapping confidence scores and cell type abundances in the reference (i.e., E1_L1) and seqFISH query datasets corresponding to Figure 2; the left dot plot shows the number of cells within cell types present in the reference and query datasets; the right dot plot shows the medians of SageNet’s confidence scores by cell type, with the width of bars corresponding to standard deviations. B. Four seqFISH query datasets in the true physical coordinates; cells are coloured by cell type with transparency according to SageNet’s mapping confidence score.

**Supplementary Figure 8.**
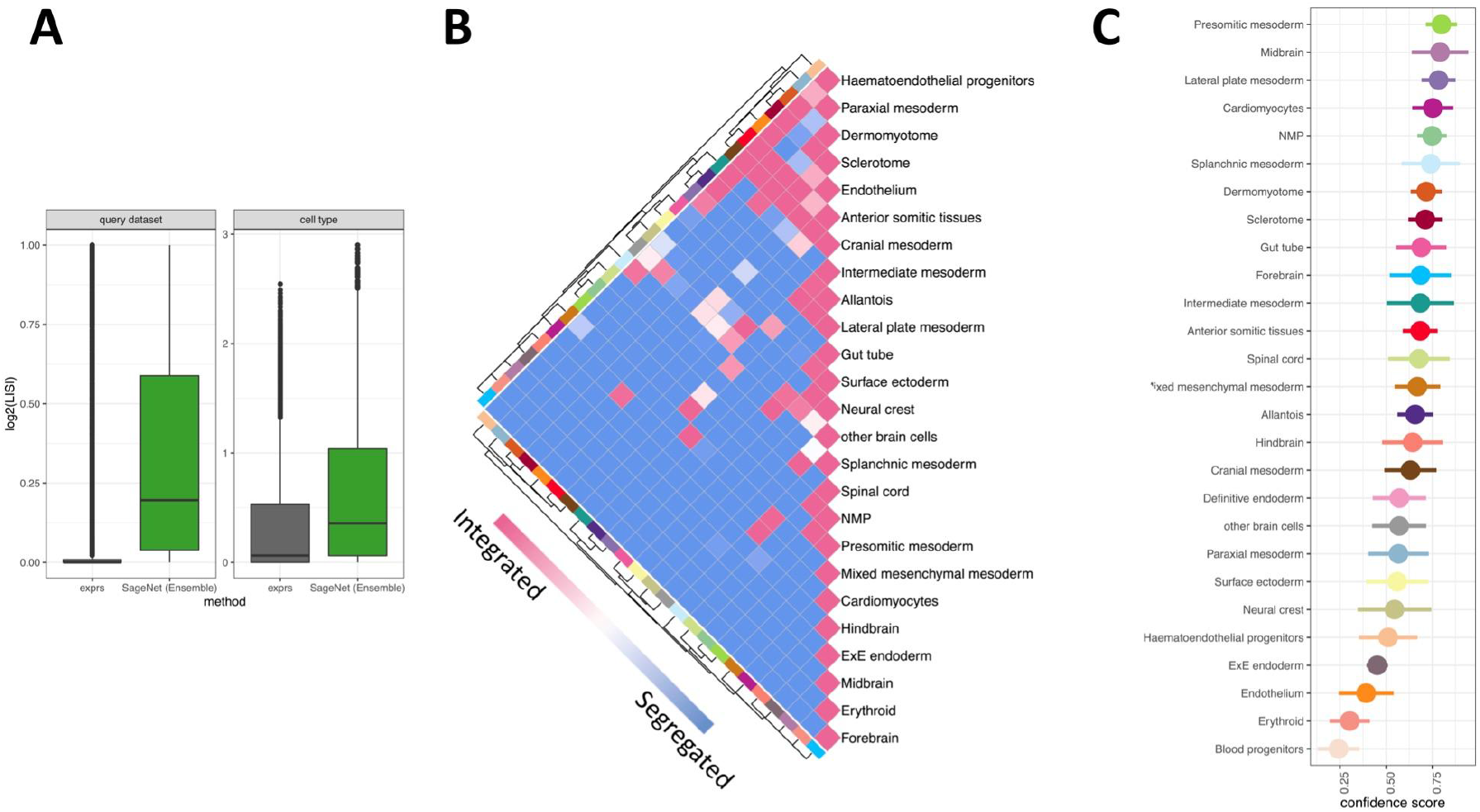
Cell type and dataset mixing, SageNet’s cell type contact map, and confidence scores on mouse seqFISH and scRNAseq embryo data. A. Boxplots of log2–LISI scores computed on the reconstructed spaces according to expression or SageNet (Ensemble). Left: query datasets as labels (scRNAseq vs. seqFISH), right: cell types as labels. B. Cell type contact maps computed on the reconstructed space shown in Figure 3B, using SageNet (Ensemble). Cell types are clustered based on the contact values by hierarchical clustering. C. Distribution of SageNet’s mapping confidence scores corresponding to Figure 3B; dot plot shows the medians of SageNet’s confidence scores by cell type, with the width of bars corresponding to standard deviations.

**Supplementary Figure 9.**
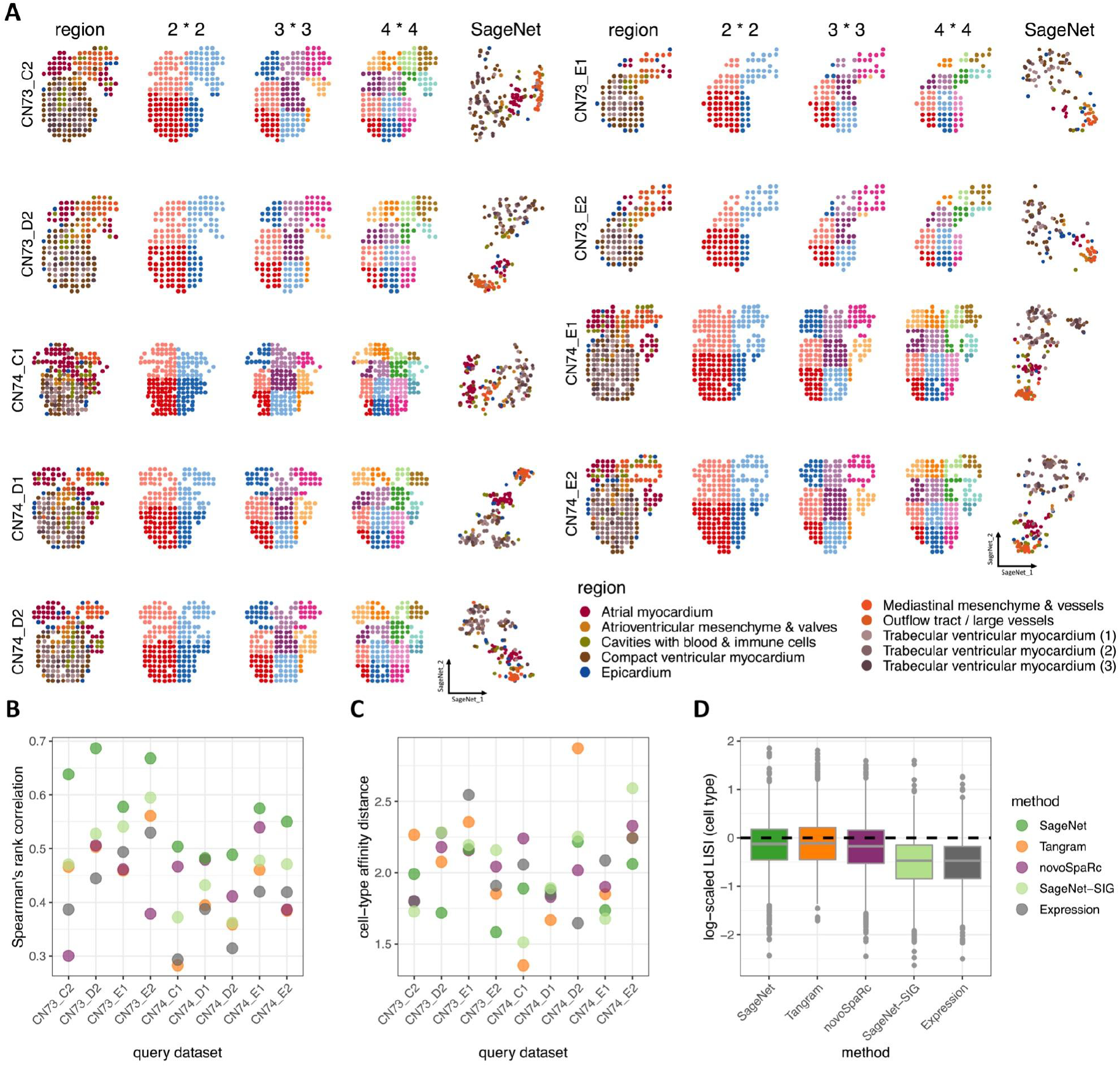
Cross-validation of developing human heart. A. Spot-based Spatial Transcriptomics (ST) samples, the partitionings, and the corresponding reconstructed spaces; each sample is shown with its spots coloured by functional region, 2×2, 3×3, and 4×4 grids and the reconstructed space using the leave-one-out mapping scheme (left to right). B. Spearman’s rank correlation between each method’s predicted cell-cell distance matrix and the true spatial distance matrix, by sample. Higher values indicate a more accurate reconstruction. C. Matrix distances between each method’s predicted region contact map and the contact map computed using the true physical coordinates, by sample. Lower values indicate a more accurate reconstruction. D. Boxplot of log-scaled LISI ratios for cell type mixing; The LISI ratio for each cell is calculated by dividing the LISI score computed for the cell in each method’s reconstructed space by the LISI score computed using the true physical space. Values closer to 0 indicate a more accurate reconstruction.

**Supplementary Figure 10.**
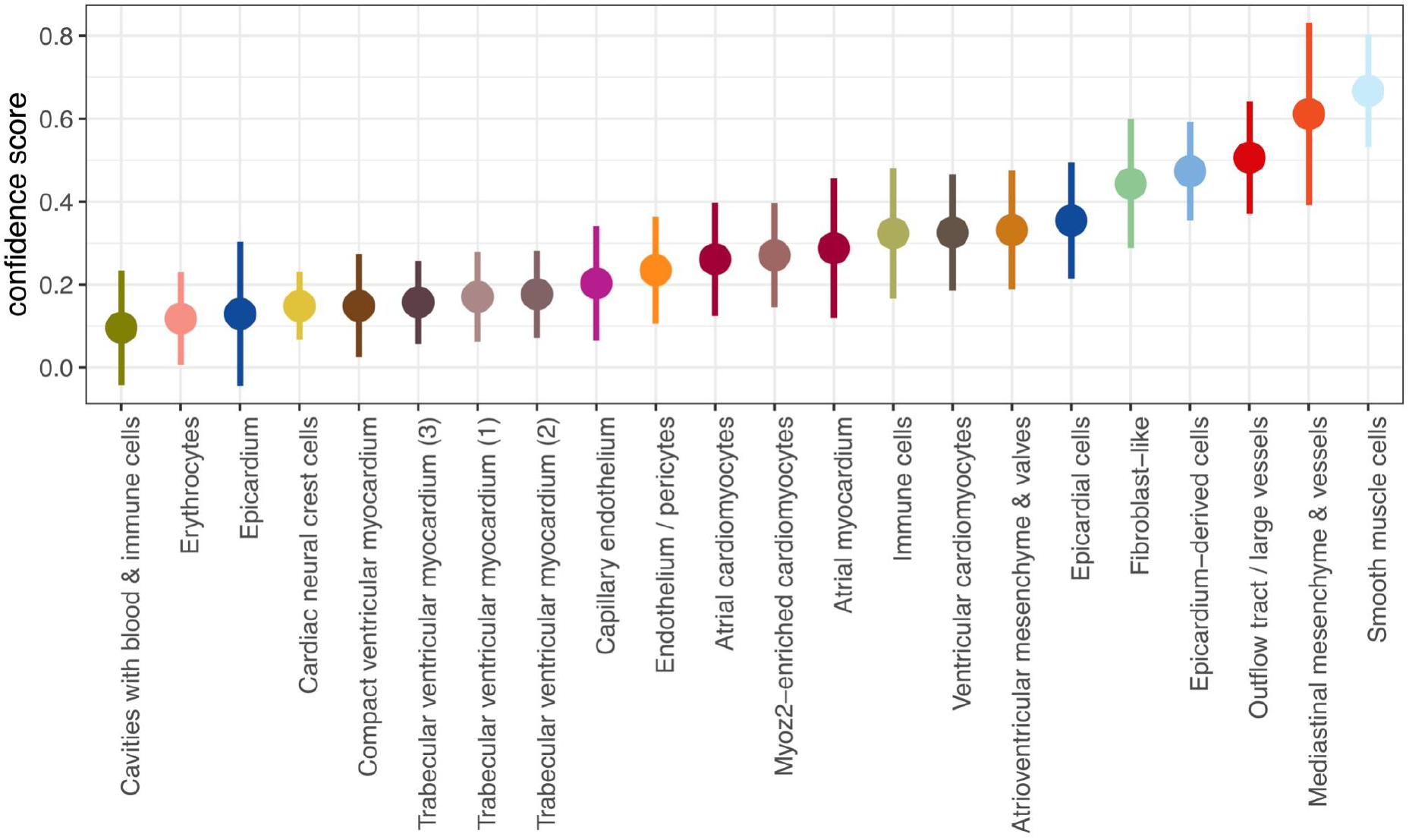
Mapping confidence of the joint embedding of ST and scRNAseq developing human heart datasets. Dot plot shows the medians of SageNet’s confidence scores by cell type, with the width of bars corresponding to standard deviations.

**Supplementary Figure 11.**
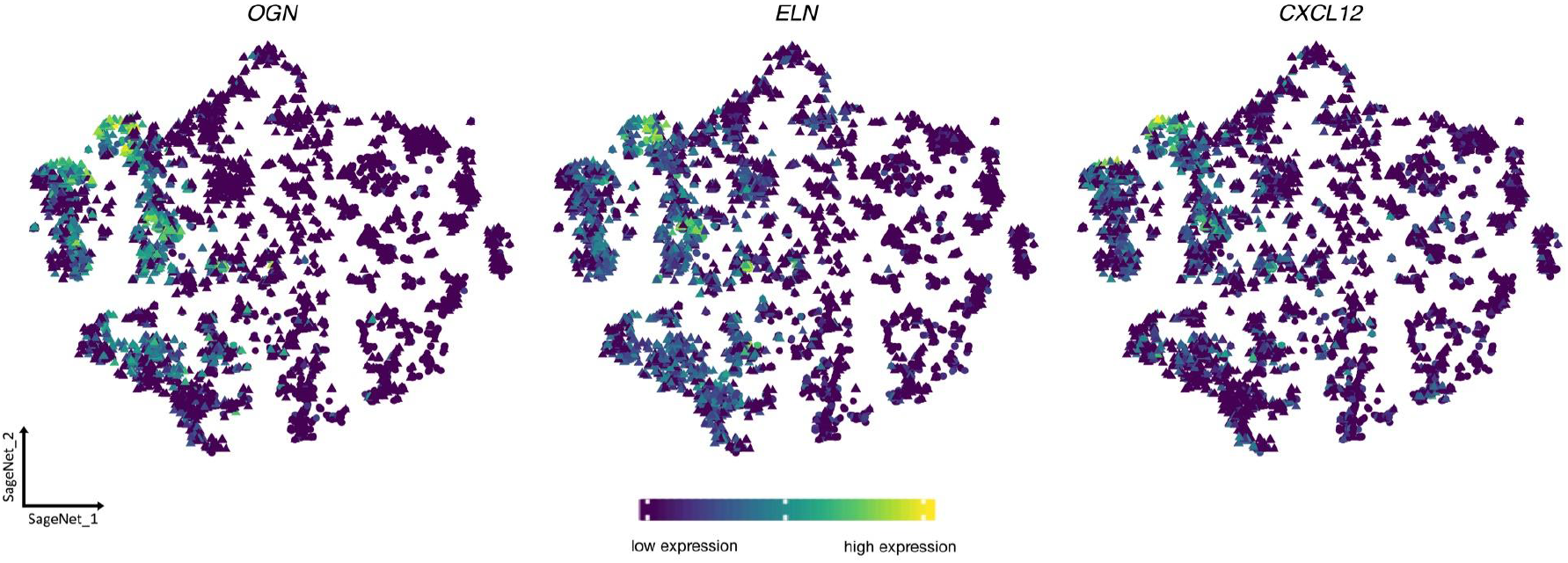
Expression of three smooth muscle cells’ markers Each panel displays gene expression of a marker in the SageNet reconstructed space as shown in Figure 4F.

**Supplementary Figure 12.**
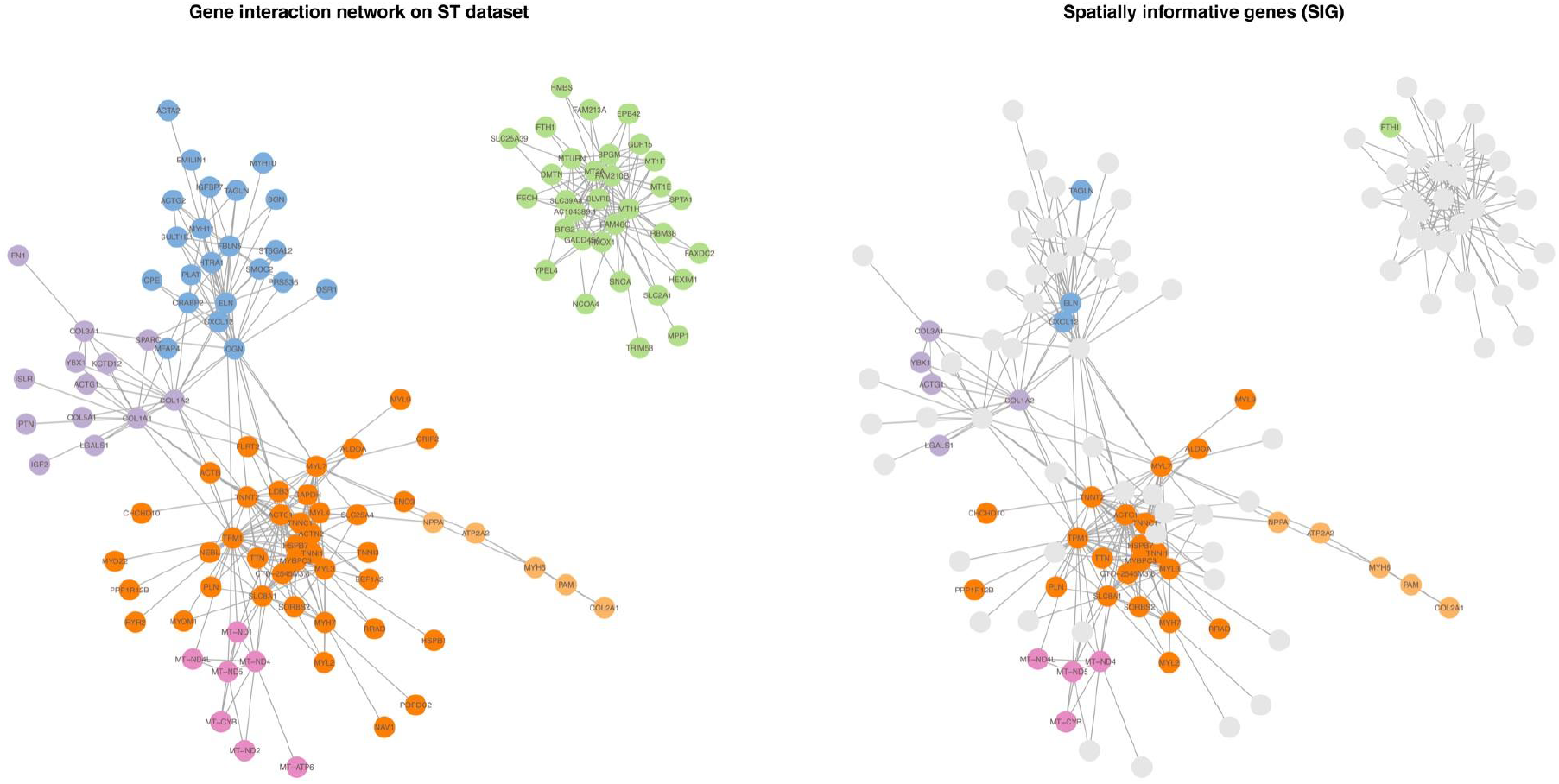
The gene interaction network on the whole developing human heart ST reference dataset (from Figure 4). The gene interaction network (GIN) built on the ST reference dataset corresponding to Figure 4; Each node represents a gene and each edge represents the statistical relevance (positive partial correlation estimated by GLASSO) of the two genes. Left: Isolated genes are removed and other genes are coloured based on their gene modules according to Leiden clustering. Right: SIGs inferred by SageNet are highlighted in the graph.

## Notes

### Competing Interest Statement

The authors have declared no competing interest.

